# ERK-mediated Curvature Feedback Regulates Branching Morphogenesis in Lung Epithelial Tissue

**DOI:** 10.1101/2021.07.11.451982

**Authors:** Tsuyoshi Hirashima, Michiyuki Matsuda

## Abstract

Intricate branching patterns emerge in internal organs because of the repetitive presence of simple deformations in epithelial tissues. During murine lung development, epithelial cells in distal tips of a single tube require fibroblast growth factor (FGF) signals generated by their surrounding mesenchyme to form repetitive tip bifurcations. However, it remains unknown how the cells employ FGF signaling to convert their behaviors to achieve the recursive branching processes. Here we show a self-sustained epithelial regulatory system during the murine lung branching morphogenesis, mediated by extracellular signal-regulated kinase (ERK), which acts as a downstream driver of FGF signaling. We found that tissue-scale curvature regulated ERK activity in the lung epithelium using two-photon live cell imaging and mechanical perturbations. ERK was activated specifically in epithelial tissues with a positive curvature, regardless of whether the change in curvature was attributable to morphogenesis or artificial perturbations. Moreover, ERK activation accelerated actin polymerization specifically at the apical side of cells, and mechanically contributed to the extension of the apical membrane, leading to a decrease in epithelial tissue curvature. These results indicate the existence of negative feedback loop between tissue curvature and ERK activity beyond scale. We confirmed that this regulation was sufficient to generate the recursive branching processes by a mathematical model. Taken together, we propose that ERK mediates the curvature feedback loop underlying the process of branching morphogenesis in developing lungs.

## Introduction

Characteristic patterns are generated in developing tubular organs through the repeated occurrence of tissue morphogenetic modes, such as elongation, bending, and bifurcation ^1–4^. During murine lung development, the complex branched architecture is stereotypically created from a simple epithelial tube, by assembling morphogenetic modes as building motifs with a spatio-temporal order ^2^. Previous genetic studies have revealed the essential molecules and signaling pathways underlying lung epithelial morphogenesis ^5–7^. Moreover, previous studies have proposed how cellular behaviors and mechanical forces determine the tube shape ^8–10^. Thus, the interplay between chemical signals and cell mechanics in constituent cells determines the organized periodic branched pattern, though it is not completely understood yet.

Several pharmacological and genetic studies have demonstrated that fibroblast growth factor (FGF) signaling is a key regulator of lung branching morphogenesis during murine development ^11^. The treatment of an isolated epithelium in which the mesenchyme is removed with different FGF ligands has clarified the distinct roles of FGFs in lung epithelial morphology. For example, FGF1 and FGF10 induce bud formation, while FGF7 gives rise to a balloon-like shape, and any of these ligands could promote epithelial proliferation ^12–14^. An investigation of the spatio-temporal expression pattern uncovered the fact that *Fgf10* is expressed locally in the mesenchyme surrounding the distal bud of the lung epithelium ^14^, and its main receptor, *Fgfr2b*, is homogeneously expressed in the lung epithelium ^12^. These results led to hypothesize that the *Fgf10* prepattern initiates branching during lung development ^14^; this was partially supported by results of previous studies on mice in which *Fgf10* or its main receptor *Fgfr2b* were knocked out and showed the complete absence of lungs ^15–17^. However, it was demonstrated that the ubiquitous overexpression of *Fgf10* in its knocked out background could restore the branching process ^18^, indicating that the *Fgf10* expression is required but the precise prepattern is dispensable for the murine lung branching morphogenesis. Thus, we explored how FGF signals are converted to cellular behaviors underlying tissue morphogenesis.

A main FGF downstream signaling pathway, the extracellular signal-regulated kinase ERK/MAP kinase pathway, includes a three-tiered kinase cascade, i.e., RAF - MEK – ERK ^11^. The genetic inactivation of either *Mek1*/*2* or *Erk1*/*2* specifically in the airway epithelium results in lung agenesis ^19^, implying that ERK and its activation were essential for lung development ^20^. Previously, immunofluorescence staining results suggested that the phosphorylation of ERK was most likely to occur in the distal tips of the lung epithelium and may, thus, be involved in the budding process during lung development ^21–23^. FGFR2-mediated ERK activation reportedly upregulates the expression of *Anxa4*, whose products are able to bind with F-actin bundles, and promote cell migration within epithelial tissues ^22^. These reports revealed the role of ERK activity in cellular behavior during lung development. However, it is still unclear how the epithelial cells collectively behave to generate the periodic branching morphogenesis mediated via ERK signaling.

In this study, we show a physical basis for self-sustained branching morphogenesis by combining two-photon live cell imaging of a Förster resonance energy transfer (FRET)-based biosensor, mechanical perturbation, and mathematical modeling. We find that ERK activation occurs specifically in epithelial tissues with a positive curvature, and promotes cell extension through actin polymerization at the apical edges, resulting in a decreased tissue curvature. Thus, our study clarifies that the negative feedback loop between the tissue curvature and ERK activity underlies the process of repetitive branching morphogenesis in developing murine lung epithelial tissues.

## Results

### 1. ERK activity is correlative to tissue curvature of lung epithelium

We first examined the impact of ERK activity on lung epithelial morphology during murine development. We cultured developing murine lungs dissected at embryonic day (E) 12.5 in the presence or absence of PD0325901, an inhibitor of the ERK activator MEK, under explant culture conditions. The growth of 1-day cultured lung epithelial tissues was markedly impaired by treatment with an MEK inhibitor (Figures 1A, 1B), indicating the importance of ERK activation for epithelial morphogenesis during this stage.

**Figure 1.**
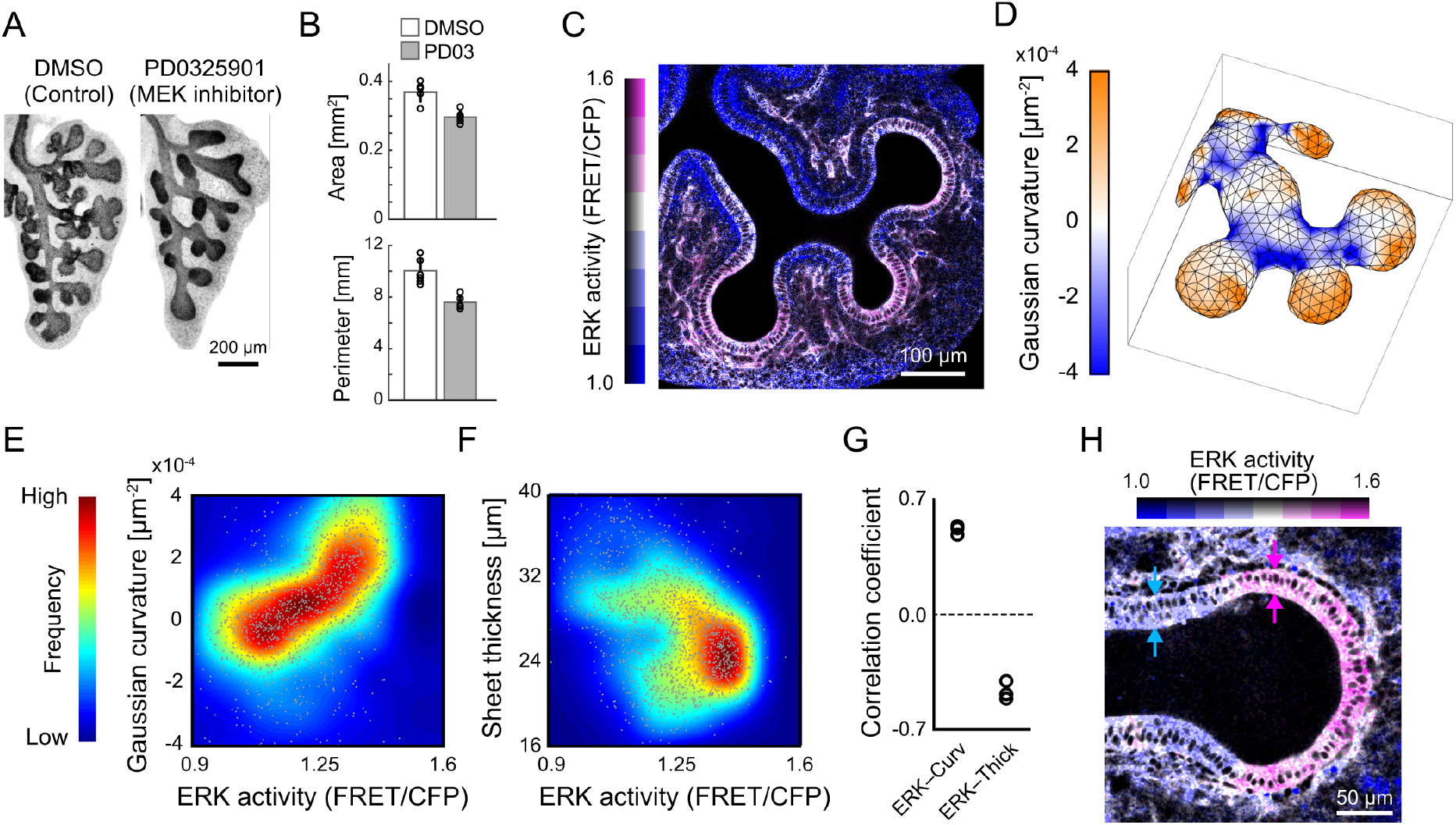
Relationship between ERK activity and tissue geometry in epithelium of intact lung lobes. (A) Immunofluorescence images of anti-E-cadherin staining in an explant culture of the murine lungs for 1 day dissected from E12.5, after treatment with dimethyl sulfoxide (DMSO) as a control (left) and 1 µM PD0325901 as a specific MEK inhibitor (right). Scale bar, 200 µm. (B) Area and perimeter of epithelial tissues from the projected images. The number of buds: 34.2±2.8 for the DMSO control and 19.2±2.8 for the PD0325901 treatment. Data represent mean and standard deviation (SD) values. N=5, two-sample t-test, p<0.001. (C) Representative ERK activity map in a developing lung at E12.5. Scale bar, 100 µm. (D) Gaussian curvature map on the basal side of the lung epithelium. (E, F) Scatter plot of ERK activity versus Gaussian curvature (E) and that of ERK activity versus sheet thickness (F). Color indicates frequency. n=1495 for (E) and 1743 for (F), N=3. (G) Correlation coefficient of (E) and (F). N=3, one-sample t-test, p=0.001 (E), and p=0.004 (F). (H) ERK activity map in the distal region of a lung dissected at E12.5. Arrows facing each other indicate the sheet thickness at the distal tip (magenta) and the distal stalk (blue). Scale bar, 50 µm.

Then, we visualized the spatial distribution of ERK activity in lung lobes dissected from murine embryos that ubiquitously expressed the FRET biosensor for the ERK activity ^24–26^ via whole tissue two-photon live imaging (Figure 1C, Movie 1). FRET images showed that the ERK activity in the epithelium was well correlated with tissue curvature; the ERK activity was higher in the curved sheet protruding the mesenchyme (convex), while it was lower in the sheet bent towards the lumen (concave) (Figure 1C). Although endothelial cells exhibited relatively high levels of ERK activity, they were less involved in epithelial branching morphogenesis ^27^; thus, we focused on ERK activity in the epithelial tissues. The basal surface of the epithelial tissues was extracted for quantification, and positions at nodes of generated meshes were embedded on the surface, enabling us to map chemical and geometrical quantities, such as ERK activity, epithelial tissue curvature, and tissue thickness (Figures 1D, S1A). As expected, there is a positive correlation between the ERK activity and tissue curvature (ρ=0.51, where ρ is the Spearman’s rank correlation coefficient; Figures 1E, 1G, S1B), which indicates that ERK activation would occur mainly in convex regions. The curvature-dependent ERK activation was observed in the developing lung epithelium under the explant culture condition (Figure S1C, Movie 2). Moreover, there is a negative correlation between the ERK activity and tissue thickness (ρ=−0.48; Figures 1F, 1G). As observed in distal buds, the epithelial tissue was thinner in the tip region, where ERK was activated, than in the stalk region, where ERK was inactivated (Figure 1H). These results suggest the existence of interplay between the ERK activity and cell morphology in developing murine lungs.

### 2. ERK activity follows morphological change of isolated epithelial tissues

To analyze how ERK activity is regulated in the lung epithelium, we isolated epithelial tissues from mesenchymal tissues, and embedded them in Matrigel with culture media containing FGFs for ex vivo culture (Figure 2A). This simplified experimental system allows us to explore epithelial response to external chemical and mechanical stimuli under regulated conditions. As reported previously ^12,13^, treatment with 500 ng mL^-1^ FGF1 for one day induced branched epithelial buds, while treatment with 100 ng mL^-1^ FGF7 gave rise to a cyst-like structure with an enlarged lumen (Figure 2B). PD0325901 treatment with FGF1 resulted in the collapse of the lumen and absence of large deformations; this was also observed with the absence of growth factors with 1-day-old culture (Figure 2B). We then examined the spatial distribution of ERK activity in epithelial tissues, one hour after chemical treatment. FGF1 activated ERK exclusively at the distal buds in a manner similar to that observed with the mesenchyme (Figure 1C), through ligand endocytosis from the basal side of cells (Figure S2, Movie 3), despite the homogeneous expression of four FGF receptors, i.e., FGFR1, FGFR2, FGFR3, and FGFR4 in the distal epithelial tissues ^12,18,28,29^. In contrast, FGF7 activated the ERK almost homogeneously in the epithelial tissues (Figure 2C). This homogeneous ERK activation was maintained during the tissue morphogenetic process (Figure S3, Movie 4). As expected, ERK activity was homogeneously low in epithelial cells treated with PD0325901 or cultured without any growth factors (Figure 2C). These results suggest that localized ERK activation at the distal tips is key to the process of branching morphogenesis.

**Figure 2.**
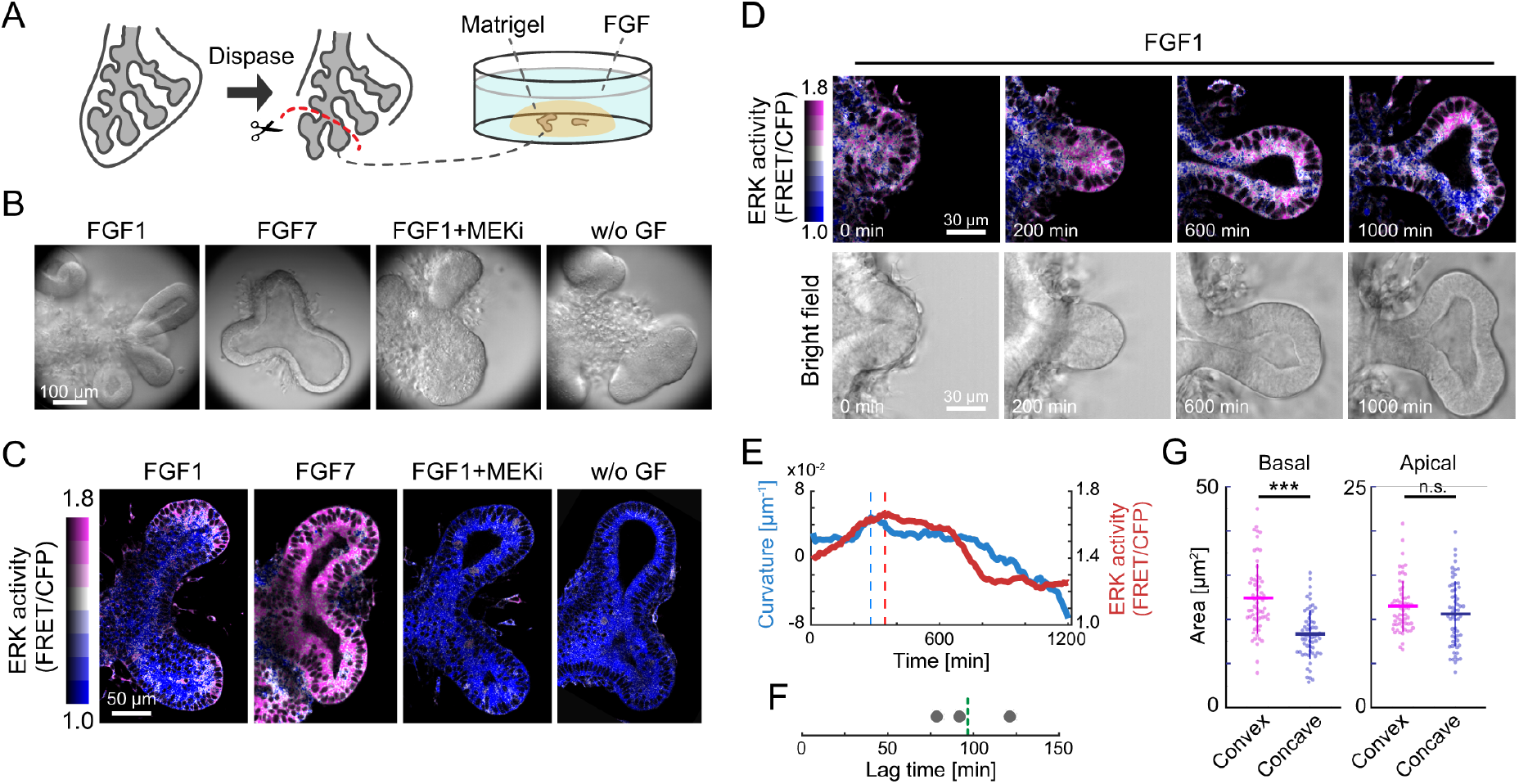
ERK activity and tissue curvature in the isolated epithelium. (A) Schematics of the culture system for the isolated lung epithelial tissues. (B) Bright field images of isolated epithelial tissues subjected chemical treatments for 1 day from E12.5. Concentrations: 500 ng mL^-1^ FGF1, 100 ng mL^-1^ FGF7, 1 µM PD0325901 (MEKi). Scale bar, 100 µm. (C) ERK activity of isolated epithelial tissues subjected to chemical treatments shown in (B) for 1 hour from E12.5. The duration of administration was so short, compared to the timescale of tissue morphogenesis, that the epithelial tissues showed similar shape in any case but different ERK activity patterns. Scale bar, 50 µm. (D) Time-lapse images of ERK activity (upper) and bright field (lower) in the case of 500 ng mL^-1^ FGF1 treatment to the the isolated lung epithelial tissue. Time origin is at 4 hours after the FGF1 treatment. Scale bar, 30 µm. (E) Tissue curvature (blue) and ERK activity (red) at the tip apex over time (E). Vertical dotted lines indicate the peak time for the tissue curvature and ERK activity, respectively. (F) Lag time from the peak for tissue curvature to the peak for ERK activity. N=3. (G) Basal area (left) and apical area (right) of individual cells in the convex and concave regions of isolated epithelial tissues. Bold and error bars represent the mean and SD values, respectively. n=60, N=3. Welch’s t test, p<0.001 (left) and p=0.18 (right).

To examine the spatio-temporal dynamics of ERK activity during the branching of isolated epithelial tissues, we performed two-photon live imaging under FGF1 supplemented conditions. We found that the ERK activity increased during the protrusion of epithelial tips, and decreased when the tip apex became flat during the tip bifurcation process (Figure 2D, Movie 5). Notably, no mesenchymal cells were observed around the distal tip of the epithelium (Figure 2D, Movie 5), suggesting that tip bifurcation could occur in a self-sustaining manner without mesenchymal cells. The quantification of the ERK activity and tissue curvature at the tip apex shows that the decrease in ERK activity follows that in curvature, with average delay of 97 min and the standard deviation of 21 min (Figures 2E, 2F). This observation allowed us to hypothesize that the ERK activity would be controlled by the curvature of the epithelial cells.

To clarify the morphological differences in cells between the convex and concave regions of epithelial tissues, we measured the basal and apical areas of epithelial cells through whole-tissue staining using E-cadherin. The basal cell area of isolated epithelial tissues in the convex region was significantly larger than that in the concave region (Figure 2G, left), while the apical cell area in the convex region was almost equivalent to that in the concave region (Figure 2G, right). The epithelial basal edges act as an interface for cells to receive growth factor signals from external environments that activate ERK (Figure S2). Therefore, we speculate that the extension of basal membranes would trigger ERK activation in the convex regions of epithelial tissues.

### 3. Morphological change of the epithelial tissues controls the ERK activity

To test whether the shape of epithelial tissues controls the ERK activity, we performed a physical perturbation assay using isolated epithelial tissues. We embedded isolated epithelial tissues within the FGF1-containing Matrigel filled inside a polydimethylsiloxane (PDMS) chamber, and uniaxially compressed it by 33% either parallelly or vertically to the distal-proximal axis of the lung epithelium (Figure 3A). Time-lapse images obtained during this process enable us to examine the ERK activity response to perturbations in different axes (Figures 3B, 3C). For parallel compression, the curved epithelial tissues at the tip apex became flattened (Figure 3D) and ERK activity decreased exclusively in the flat region (Figure 3E, Movie 6). However, for vertical compression, the tip apex of the epithelium became sharp (Figure 3D) and ERK activation followed the change in tissue curvature (Figure 3E, Movie 7).

**Figure 3.**
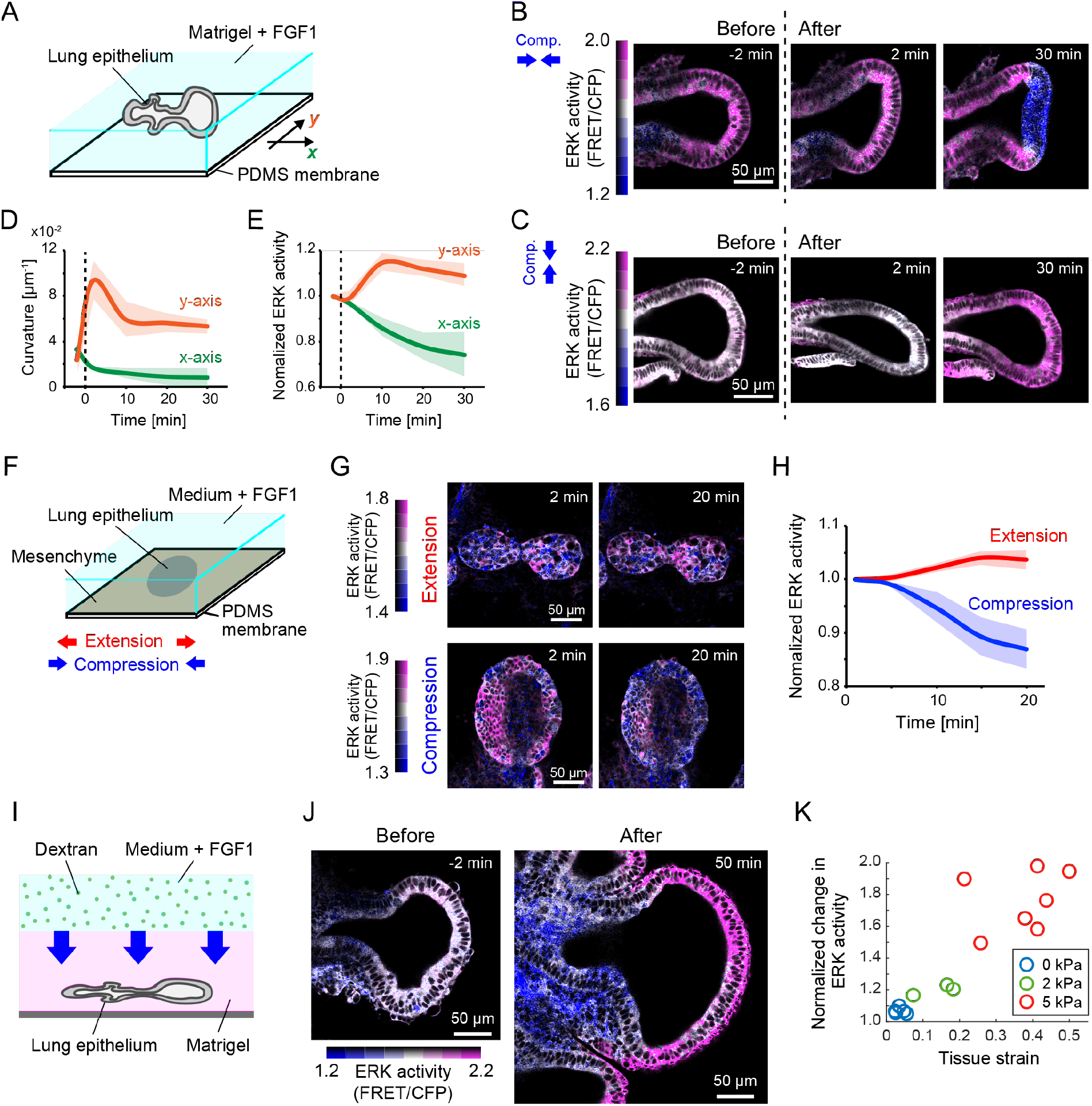
Response of ERK activity to mechanical perturbations. (A) Schematics of PDMS-based uniaxial compression assay performed on an isolated epithelium under the 3D culture condition. The *x* and *y* axes indicate directions parallel and vertical to the proximo-distal axes of the epithelium, respectively. (B, C) Time-lapse images of the ERK activity for parallel compression (B) and vertical compression (C). Time origin indicates the timing of mechanical compression. Scale bars, 50 µm. (D, E) Tissue curvature at the tip apex (D) and normalized ERK activity (E) over time. Colors represent the axes of compression, as shown in (A). Data represent mean and SD values. N=3. (F) Schematics of PDMS-based uniaxial mechanical perturbations under the 2D culture condition. (G) Time-lapse images of the ERK activity in the isolated lung epithelium for the extension (upper) and compression (lower). Time origin indicates the timing of mechanical perturbations. Scale bar, 50 µm. (H) Normalized ERK activity for the extension (red) and the compression (blue) processes over time. Data represent mean and SD values. N=3. (I) Schematics of osmolyte-based uniaxial compression assay from the top under the 3D culture condition. Blue arrows represent the displacement of interface between the Matrigel and the osmolyte-containing culture medium due to the gel shrinkage. (J) Snapshots of the ERK activity in the isolated lung epithelium before and after the osmolyte-based compression (lower). Scale bar, 50 µm. (K) Normalized ERK activity versus the tissue strain in the distal tip. The data was obtained at 50 min after the dextran treatment. N=5, 3, 7 for 0, 2, 5 kPa, respectively.

We further examined the ERK activity response to cell deformation. The manipulation of cell shape in curved epithelial tissues remains unfeasible under 3D culture conditions; thus, we performed a strain perturbation assay under planar culture conditions using dissociated lung epithelial cells (Figure 3F). Under planar culture conditions, the stretching of cells led to ERK activation, while their compression led to ERK inactivation (Figure 3G, 3H). In addition, we performed osmolyte-based compression assay^30^, in which the Matrigel shrinks due to the difference in osmotic pressure with the culture medium (Figure 3I). Treatment with dextran molecules as the osmolyte led to flattening of the lung epithelial tissues by the global compression from the top (Movie 8). We found that ERK was significantly increased in the tips, which of the epithelium were extended on the plane orthogonal to the compression axis (Figure 3J). Remarkably, the ERK activation exhibited a positive correlation with the level of tissue extension in the tips (Figure 3K). Together, these results indicate that the strain on the cell area is key to curvature-dependent ERK activity regulation. Notably, the surface area of the basal side of cells was significantly larger in the convex region with high ERK activity than in the concave region with lower ERK activity (Figure 2G). Together, epithelial tissue curvature controls the ERK activity in cells, most likely through the strained state of their basal membranes.

### 4. ERK activation determines the epithelial cell shape via actin polymerization at the apical side

Next, we investigate how the ERK activity contributes to epithelial morphology and mechanics. First, we observed the change in the cell shape of isolated epithelial tissues treated with the MEK inhibitor PD0325901. Drug treatment led to an inhibition in the ERK activity within 30 min after administration (Figure S4A), and the apico-basal length of epithelial cells became longer than that observed in the control within 2 hours (Figure 4A, Movie 9). Hereafter, we refer to the apico-basal length of cells as the cell height, and the cell length orthogonal to the apico-basal axis as the cell width (Figure 4B). ERK inactivation in cells treated with PD0325901 lengthened the cell height to 124% and shortened the cell width to 92% on an average, despite no significant changes in the cell size, compared to mock-treated cells (Figure 4C). This trend is consistent with that observed in the dissected murine lungs (Figure 1E–G). These results indicate that the ERK activity regulates the cell height, while maintaining the cell size.

**Figure 4.**
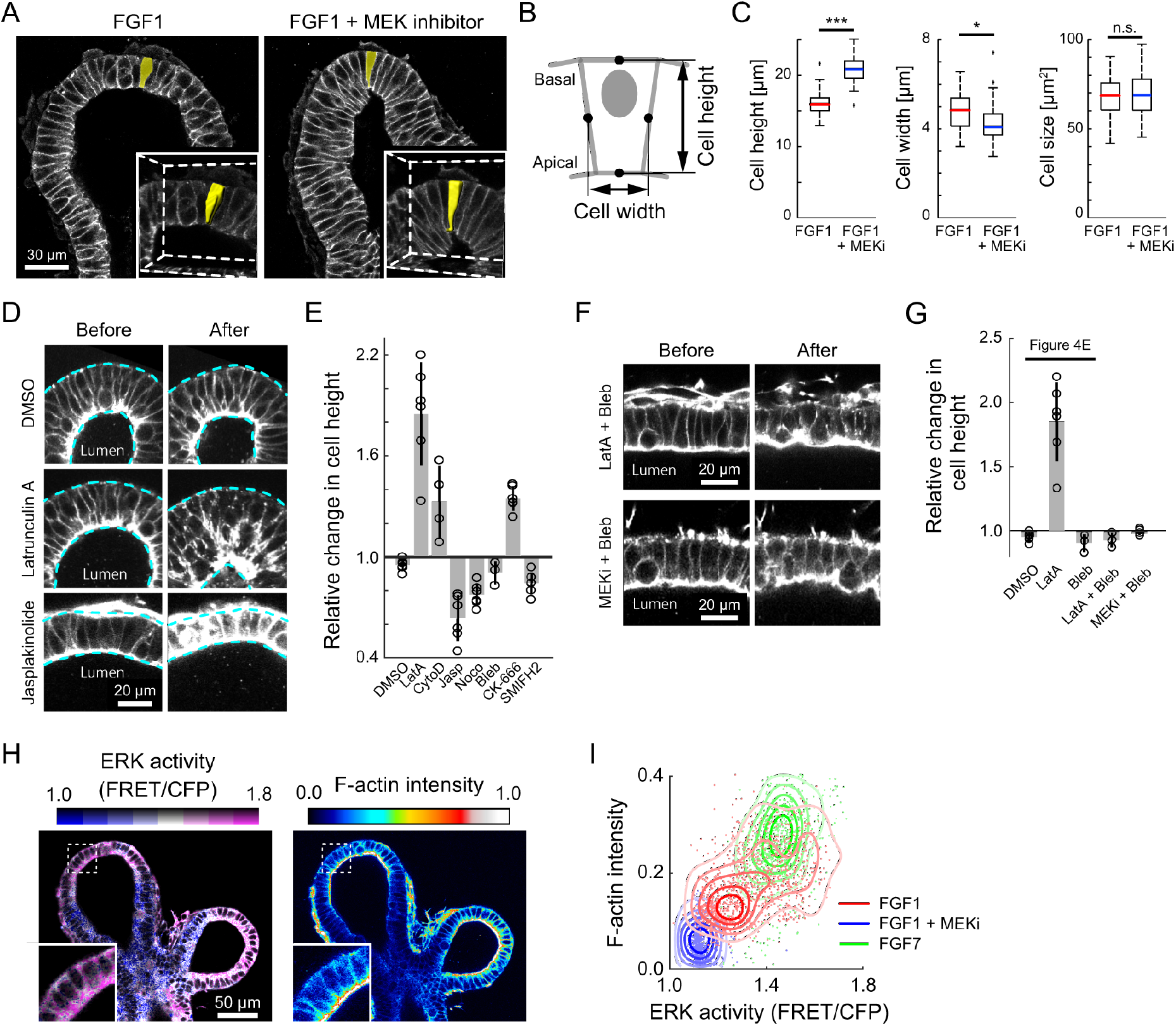
Response of cell morphology to chemical perturbations. (A) Change in the cell shape of an isolated epithelium upon treatment with either 500 ng mL^-1^ FGF1 (left) or the mixture of 500 ng mL^-1^ FGF1 and 1 µM PD0325901. The whole-mount immunofluorescence of anti-E-cadherin was used for the visualization. The inset windows represent the magnified 3D rendering focusing on a representative single cell marked in yellow. Scale bar, 30 µm. (B) Schematics for cellular morphological measurement. (C) Quantification of cell morphological parameters, including cell height, width, and size. n=60, N=3. Welch’s t test, p<0.001 (left), p=0.013 (middle), and p=0.4 (right). (D) Change in the cell morphology of an isolated epithelium upon treatment with either DMSO control, 1 µM Latrunculin A, or 1 µM Jasplakinolide. SiR-Actin, a F-actin labeling probe, was used for visualization. Scale bar, 20 µm. (E) Relative change in the cell height upon the inhibitor treatment for cytoskeletons. Concentrations of inhibitors: 1 µM Latrunculin A, 1 µM Cytochalasin D, 1 µM Nocodazole, 30 µM Blebbistatin, 1 µM Jasplakinolide, 100 µM CK-666, and 10 µM SMIFH2. The bar graph and error bars represent mean and SD values. N≥3 for each treatment. (F) Snapshots of the cell morphology before and 40 min after the treatment with either the mixture of 1 µM Latrunculin A and 30 µM Blebbistatin or the mixture of 1 µM PD0325901 (MEK inhibitor) and 30 µM Blebbistatin. The SiR-Actin was used for visualization. Scale bars, 20 µm. (G) Relative change in the cell height upon the inhibitor treatment for cytoskeletons. The three on the left are the same as (E) and shown for a comparison purpose. The bar graph and error bars represent mean and SD values. N=3 and 4 for LatA+Bleb and MEKi+Bleb, respectively. (H) Spatial map of ERK activity and F-actin accumulation level visualized by SiR-Actin labeling upon treatment with 500 ng mL^-1^ FGF1. Inset windows show magnified views that focus on their subcellular localization. Scale bar, 50 µm. (I) Scatter plot and contour map of ERK activity in the cytosol and F-actin intensity at the apical edge. Colors indicate the administration of different chemicals; red: 500 ng mL^-1^ FGF1, n=1204, N=4, blue: 500 ng mL^-1^ FGF1 and 1 µM PD0325901 (MEKi), n=687, N=3, green: 100 ng mL^-1^ FGF7, n=787, N=3.

To explore what causes the cell shape to change mechanically, we performed inhibitor assays on cytoskeletons, such as actin, microtubule, and non-muscle myosin, which allowed us to determine the mechanical contributions of targeted cytoskeletons to the cell morphology. As inferred by a theoretical analysis (see Materials and Methods IIa), the cell height and the cell width showed an inverse relationship in the phase before cytoskeletal collapse, although it failed in the later phase owing to actin polymerization inhibition (Figure S4B). In addition, the cell width is not the parameter that can be measured robustly due to technical reasons of live imaging. Thus, we focused on the change in the cell height as a proxy for cytoskeletal perturbations. The cell height was increased markedly upon treatment with Latrunculin A or Cytochalasin D, each an inhibitor of actin polymerization, as observed in inhibiting ERK activity with PD0325901; conversely, the cell height was decreased by treatment with Jasplakinolide, which promotes actin polymerization (Figures 4D, 4E). Because F-actin remarkably accumulated at the apical side of cells (Figure S4C), we speculated that actin polymerization at the apical edge would contribute to cell width extension by pushing the neighboring cells ^31^, leading to the shortening of the cell height. To further analyze the mode of actin polymerization, we used CK-666, an Arp2/3 complex inhibitor, and SMIFH2, an inhibitor of the formin homology 2 domain. We found that the cell height was lengthened by CK-666 treatment, but slightly shortened by SMIFH2 treatment (Figure 4E), suggesting that actin polymerization would depend on the Arp2/3-mediated nucleation of branched filaments, rather than formin-mediated actin nucleation.

The cell height was also shortened by interfering with microtubule polymerization with Nocodazole; combined treatment with Nocodazole and Latrunculin A resulted in significant cell height extension via the rupture of lateral cell edges (Figures S4D, S4E). Microtubules within the lung epithelial cells are distributed along the apico-basal axis (Figure S4F), and would thus stabilize the cytoskeletal architecture cooperatively with F-actin. The inhibition of the activity of non-muscle myosin II with Blebbistatin resulted in no significant difference in cell height, compared to that observed with DMSO control (Figure 4E). Active myosin localized at the apical edges in the tips of lung epithelium despite a weak level of accumulation (Figure S4G-G”). Moreover, Blebbistatin treatment suppressed the cell elongation triggered by Latrunculin A (Figures 4F, 4G). Taken together, these results indicate that contractile squeezing of apical edges drives apico-basal cell elongation only under the inhibitory situation of actin polymerization. Importantly, treatment with the combination of Blebbistatin and PD0325901 resulted in suppression of the cell elongation as in the case of combined treatment with Blebbistatin and Latrunculin A (Figures 4F, 4G).

Because the inhibition of ERK activity and actin polymerization resulted in similar effects on the cell height, we speculated that ERK activation promoted F-actin accumulation within cells. Simultaneous imaging for the two factors revealed that F-actin accumulated at the apical side of cells exclusively in the tips where ERK activation occurred, under FGF1-treated conditions (Figure 4H). Mapping between the ERK activity and F-actin accumulation level via 3D whole-tissue imaging clearly showed a positive correlation with FGF1 (ρ=0.51), although the inhibition of the ERK activity led to a significant decrease in the F-actin accumulation at the apical side of cells throughout the epithelial tissues, and relatively uniform distribution (Figure 4I). In addition, treatment with FGF7, which activates ERK globally in the isolated epithelial tissues regardless of the tissue curvature (Figures 2C, S3), led to an increase in the apical F-actin accumulation overall throughout the epithelial tissues (Figure 4I). Thus, we found that ERK activation promotes actin polymerization at the apical edges of cells, and ultimately controls the cellular shape in the epithelial monolayer.

### 5. ERK activation flattens the epithelium via the apical actin polymerization

How does the ERK-mediated actin polymerization at the apical side of cells result in morphological changes in monolayer tissues? To answer this, we used a simple geometrical model of a two-dimensional monolayer tissue, composed of cells with identical morphologies. In this model, individual cells are represented by tetragons, the edges of which each correspond to the apical, basal, and lateral sides of cells. Then, the sheet curvature *κ* can be represented as a function of the length of these edges; especially, a change in the curvature with respect to the apical edge length *a* can be derived as ∂*k*/∂*a* < 0. Thus, the sheet curvature decreases with an expansion in the apical edge (see Materials and Methods IIb). This theoretical analysis predicts that ERK activation would decrease the tissue curvature via actin polymerization at the apical edges.

To confirm this prediction, we examined the change in epithelial tissue curvature in response to acute ERK activation. For this purpose, an isolated lung epithelium was incubated in the culture medium without any growth factors. Then, FGF1 was added into the medium to simulate ERK. The acute ERK activation promoted F-actin accumulation exclusively at the apical side of cells in epithelial distal tips; this was followed by the flattening of the epithelial sheet (Figures 5A–D, Movie 10). Moreover, the simultaneous administration of FGF1 and the MEK inhibitor PD0325901 resulted in neither F-actin accumulation nor tip flattening; this was also observed for the solvent control treatment (Figure 5B–D). These results indicate that ERK activation flattens the epithelial tissues in the tips via actin polymerization at the apical sides of cells.

**Figure 5.**
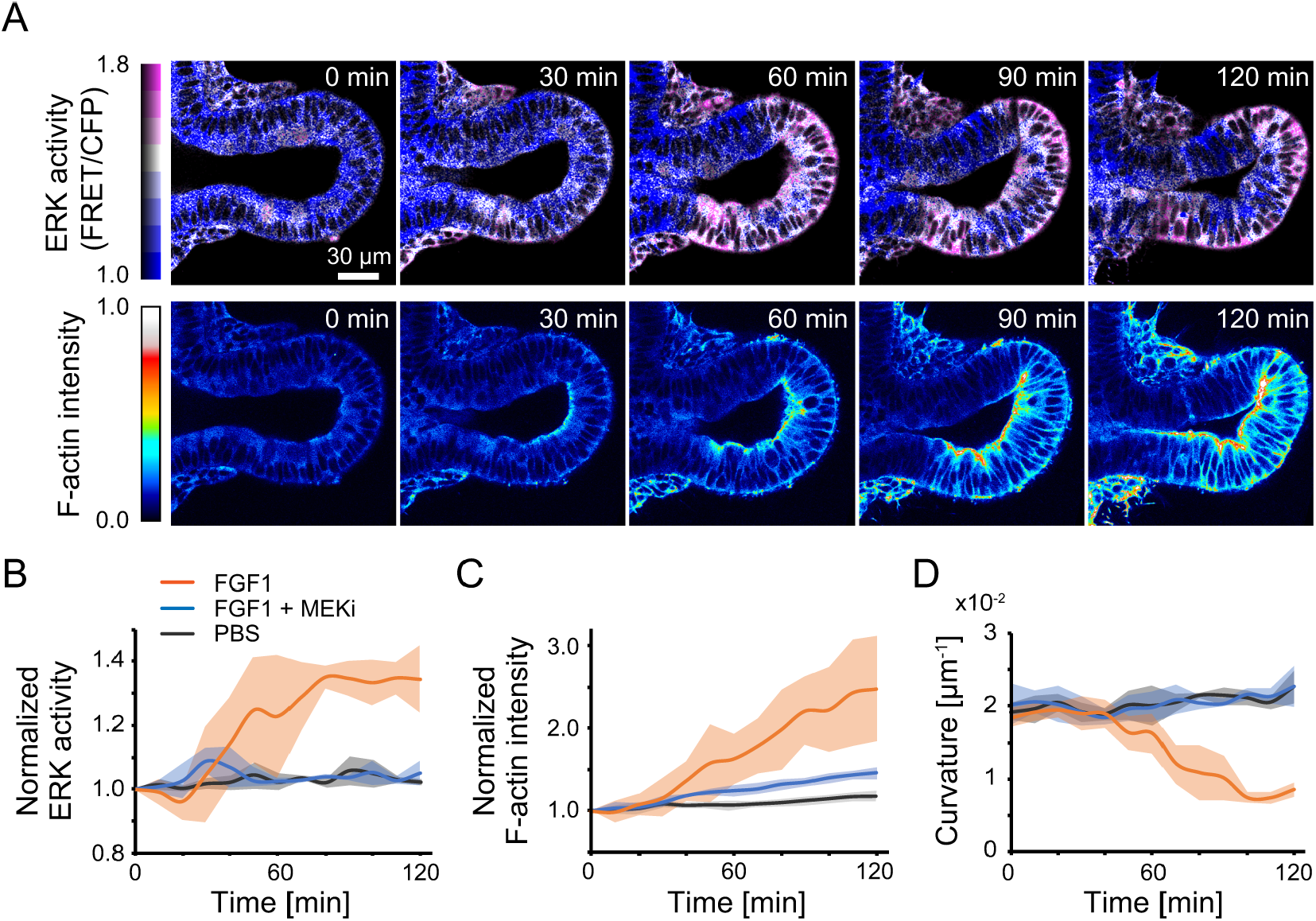
Simultaneous imaging of FGF-triggered ERK activation, actin polymerization, and tissue flattening. (A) Time-lapse images of ERK activity (upper) and F-actin intensity (lower). SiR-Actin was pretreated for F-actin visualization, 1 hour before the addition of FGF1. The time origin indicates the timing of FGF1 administration. Scale bar, 30 µm. (B–D) Normalized ERK activity in the tips (B), normalized F-actin intensity in the tips (C), and tissue curvature at the tip apex (D) over time. Colors indicate different chemical treatments, including with 500 ng mL^-1^ FGF1 and 1 µM PD0325901 (MEKi). The time origin indicates the timing of chemical administration. Data represent mean and SD values. N=4 for (B) and N=3 for (C, D).

### 6. Feedback between ERK activity and tissue curvature explains repetitive branching

Finally, we performed numerical analysis using vertex dynamics model, a cell-based mechanical model ^32–34^ to examine whether the curvature-dependent apical extension is able to produce the periodic branching pattern. The model represents epithelial cells as polygons with vertices and edges, and the edges on the apical side actively deform via F-actin dynamics that depend on the tissue-scale curvature and its decay (as described in Materials and Methods IIc). We conducted numerical simulations with an initial shape and bud diameters obtained from the experiment (Figure 6A), and examined the impact of curvature on the change in apical edge length through a model parameter *α*, using the standardized parameter set (Figure S5A). The apical edge extends/shrinks in accordance with the positive/negative values of *α* in a curvature-dependent manner (Figure 6B). Clearly, the curvature and cell height at the tip apex decreased with an increase in *α* at equilibrium (Figures 6B, S5B). Moreover, during curvature-dependent apical extension (*α* = 1), the curvature decreases at the tip apex (arrowhead, Figure 6C), but increases at both sides over time (arrows, Figure 6C), indicating that the curvature-dependent apical extension can create new buds for terminal bifurcation. This does not happen in the opposite case (*α* = −1), suggesting that curvature-dependent apical constriction is not likely to drive repetitive branching, contradictory to a previous report ^35^.

**Figure 6.**
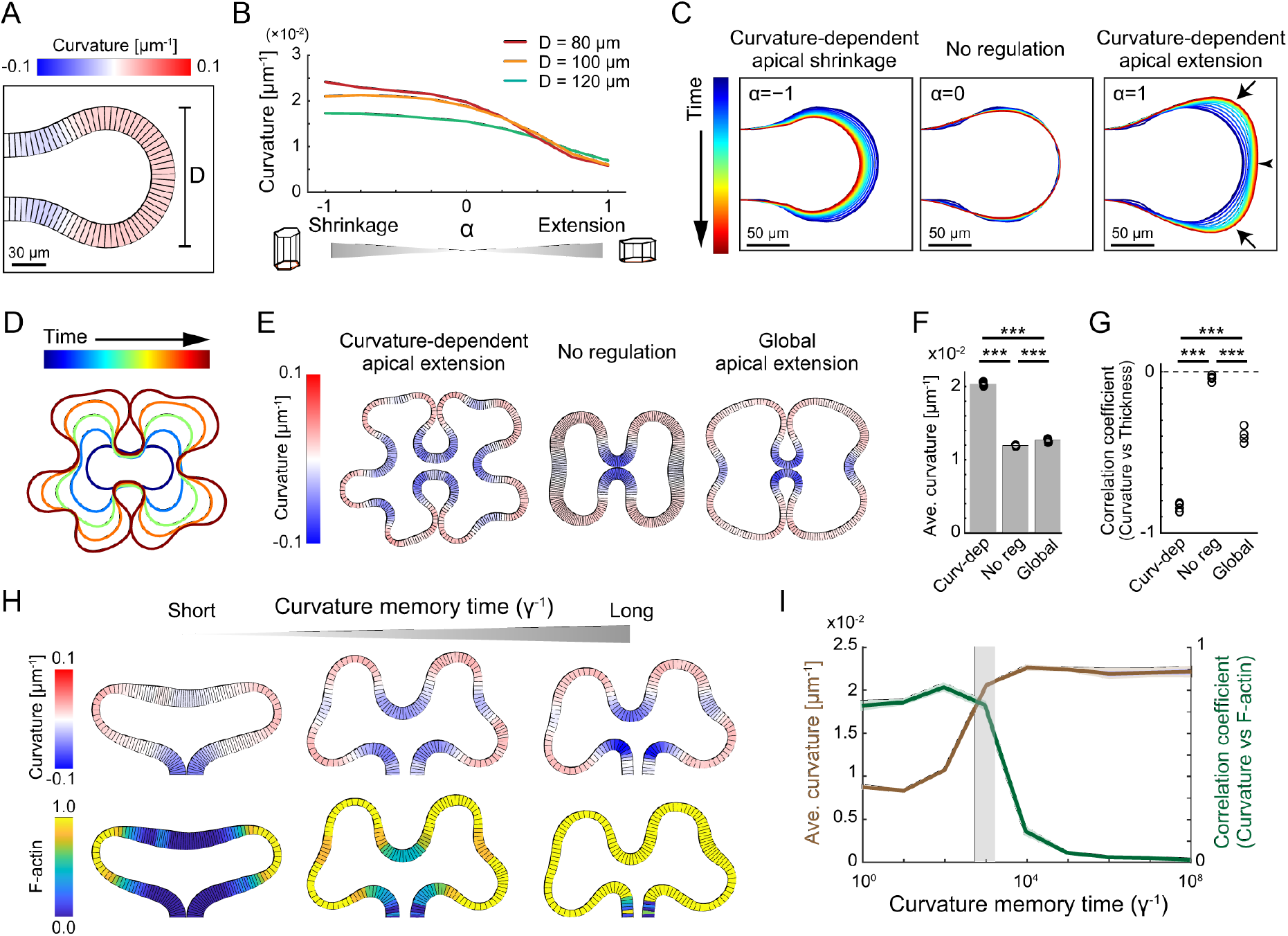
Simulation and analysis of the mathematical model. (A) Distal tip of a virtual epithelial monolayer tissue during the initial configuration of the simulation. Colors indicate the tissue curvature. D represents the diameter of the tip. Scale bar, 30 µm. (B) Tissue curvature at the tip apex over the parameter *α*. Colors indicate the tip diameter. (C) Time evolution of the basal sides of cells in different *α*. Colors indicate the time periods in simulations. The arrowhead indicates the tip apex and arrows indicate the regions with a growing curvature. Scale bars, 50 µm. (D) Time evolution of the basal sides of cells in the simulation with cell proliferation at *α*=1. (E) Generated morphology of epithelial monolayer tissues in different regimes. Curvature-dependent apical extension: *α*=1, β=0; no regulation: *α*=0, β=0; global apical extension: *α*=0, β=0.2. (F, G) Averaged curvature (F) and correlation coefficient between the tissue curvature and tissue thickness (G) for the 3 regulatory regimes. Bars represent mean values. N=5. Welch’s t-test, p<0.001. (H) Spatial maps of the curvature (upper) and F-actin level (lower) for different curvature memory time periods γ^-1^. left: γ=1, middle: γ= 10^−3^, right: γ= 10^−6^. (I) Tissue curvature (green) and correlation coefficient between the curvature and the F-actin level. Data represent mean and SD values. N=5. The gray shaded region indicates the region in which both the values are high.

We then enabled cell proliferation to occur in the model system, in order to perform in silico experiments in virtual growing monolayer tissues. The simulation of the regime of curvature-dependent apical extension and cell proliferation clearly demonstrates the formation of repetitive branching patterns (Figure 6D, Movie 11). In contrast, the repetitive branching pattern is not generated in counterpart regimes, such as no regulation on the apical edge, and global apical extension as observed for the FGF7 treatment – apical extension happens in all cells, irrespective of curvature (Figure 6E). In these counterpart regimes, there are significant differences in both the averaged curvature, a measure of curvature averaged over the entire tissue, and the correlation coefficient between the curvature and monolayer thickness (Figures 6F, 6G). Moreover, the virtual experiments show that luminal pressure, which has been proposed to be a physical factor for lung morphogenesis ^36,37^, is not essential for branching pattern formation; rather, it suppresses its constant large magnitude value (Figure S5C). These results support the fact that the exclusive occurrence of apical edge extension in cells with high curvatures drives repetitive terminal bifurcation in the growing epithelial monolayer in a self-sustaining manner.

Furthermore, numerical investigations have clarified that the retention time of curvature memory in F-actin, i.e., the inverse of the F-actin decay rate γ, is crucial for generating the branching pattern (Figure 6H, Movie 12). The averaged curvature becomes larger for a longer curvature memory, and the correlation between the curvature and F-actin becomes higher for a shorter curvature memory. Thus, there is an optimal range for the retention time of the curvature memory in F-actin (Figure 6I). This result predicts the importance of the tissue curvature hysteresis of cellular force generation via F-actin, for the emergence of repetitive branch patterns.

## Discussion

We have demonstrated a regulatory mechanism by which epithelial monolayer cells undergo repetitive branching morphogenesis through curvature sensing and ERK-mediated mechanical force generation in a self-sustained manner. Our two-photon live imaging analysis has shown that the ERK activity changes following a dynamic change in the tissue curvature, and ERK activation occurs preferentially in the curved region of epithelial tissues. Moreover, we found that ERK-mediated actin polymerization contributes to the lateral extension of the apical side of cells. Finally, we demonstrated its importance for achieving a decrease in tissue curvature during terminal bifurcation via mathematical model analyses. Thus, there is a closed negative feedback loop between the tissue curvature and ERK activity across multiple scales in the developing lung epithelial tissues (Figure 7).

**Figure 7.**
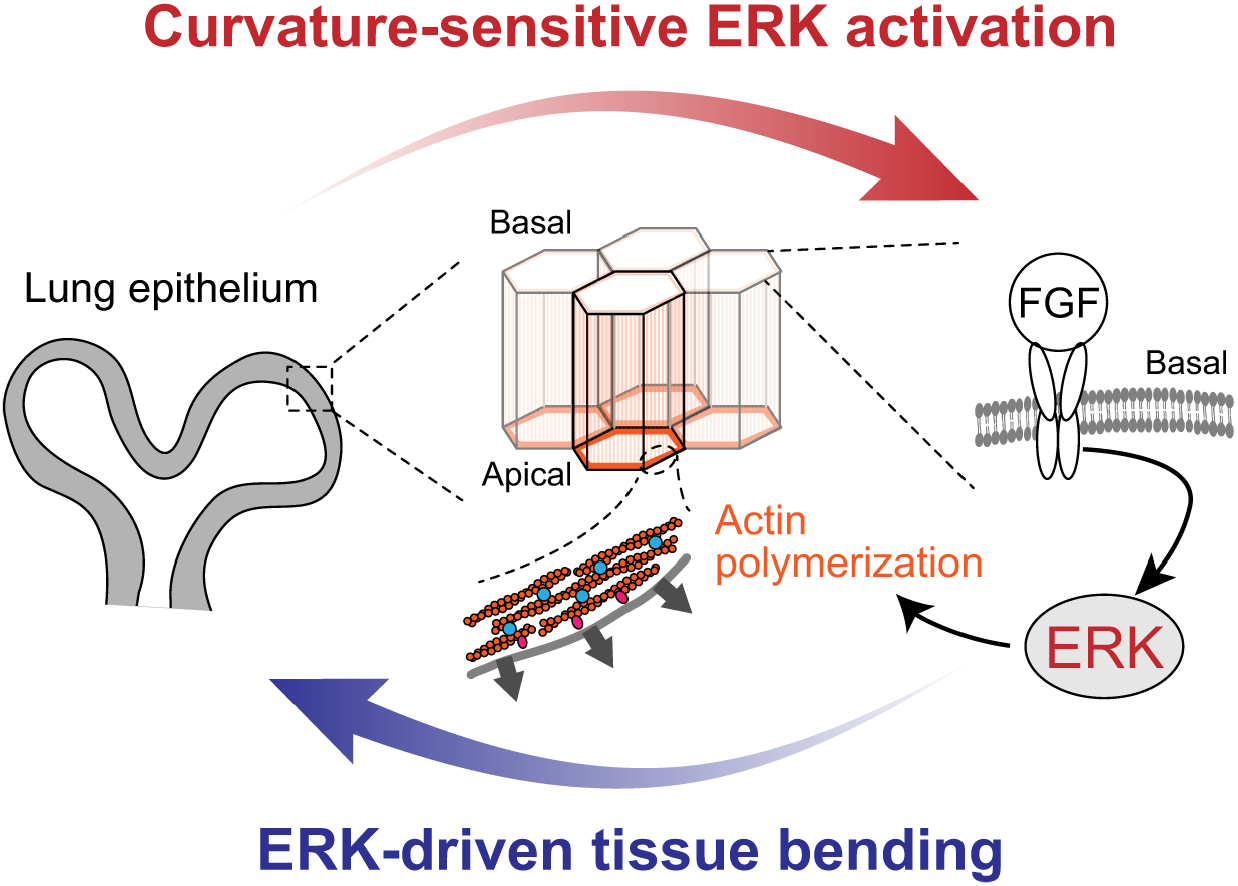
ERK-mediated curvature feedback regulates branching morphogenesis in murine lung epithelial tissue. The epithelial cells activate ERK specifically in tissues with a positive curvature via reception of extracellular FGF ligands (red arrow). ERK-mediated actin polymerization contributes to the extension of the apical side of cells, which decreases the tissue curvature (blue arrow). Thus, there is a closed negative feedback loop between tissue curvature and ERK activity across multiple scales in developing epithelial lung tissues.

The results presented in this study provide a better understanding of lung epithelial branching, in conjunction with previously determined theory-based mechanisms, based on mechanical and chemical aspects. From a mechanical viewpoint, it has been proposed that the spatial patterning of the isolated lung epithelium arises from the proliferation-induced physical instability of the epithelial layer, known as mechanical buckling ^38^, in various developing epithelial tissues ^1,39,40^. This purely mechanical model may explain the formation of an initial branching pattern under the assumption that the epithelial tissue acts as an elastic layer, but may require the physical conditions under which mechanical buckling occurs. However, our results have demonstrated that repetitive branching can be generated even at a range where proliferation-induced buckling does not occur (Figure 6E–G). We emphasize that the regime presented here is not mutually exclusive to buckling-induced patterning. Thus, ERK-mediated tissue curvature feedback results in a robust regulatory system underlying branching morphogenesis, together with the buckling regime. In addition, a simple reaction–diffusion model of substrate–depletion has been proposed for integrating chemical signals and tissue growth ^41,42^; the model assumes that lung epithelial cells consume FGF ligands that have slow diffusion rates, and thereby predicts that a concentration gradient of FGF should be generated in the extracellular matrix. However, our live imaging analysis results have shown that no FGF gradient was formed in the Matrigel, while FGF was internalized in the epithelial cells through endocytosis (Figure S2). Other reaction-diffusion systems have focused on genetic regulatory networks that can achieve the localization of *Fgf10* expression observed in vivo ^43–45^. Although the spatial distribution of the chemoattractant is dispensable for epithelial branching ^18^ (Figure 2B–D), the incorporation of such fundamental biochemical signals into our model would be useful for studying detailed systems for lung branching and patterning in vivo.

Our findings highlight the curvature-sensitive ERK activation of lung epithelial cells and raise an issue on how cells sense the tissue curvature and transmit FGF–ERK signals. Because the ERK activity response occurred within 10 min after the perturbation in the tissue curvature (Figures 3B–E), the chemical feedbacks via gene expression, e.g., the interplay between the Fgfs and Sonic hedgehog expressed in the epithelium, are unlikely. Rather, we surmise that this process involves unknown protein–protein interaction(s). We determined that the phenomenon of curvature-sensitive ERK activation in lung epithelial tissues was attributable to the strain of cell membrane or the level of membrane tension (Figures 2G, 3G–K), as shown previously for other tissues ^46–48^. Moreover, an extension of the membrane would trigger receptor-mediated endocytosis, as the FGF internalization occurred exclusively in the epithelial tissues with positive curvature (Figure S2). Multiple intracellular events, including the endocytic trafficking of FGFR and molecular sensing of plasma membrane curvature, would be involved in this process ^49–51^.

This study has focused on the core mechanism to minimally generate the self-sustained repetitive branching patterns in epithelium; however, there are other additional contributions that could achieve the robust tissue morphogenesis in vivo. For example, smooth muscle cells are not necessary for the initiation of cleft formation (Figure 2D)^52^, but should mechanically assist the cleft formation from the outside of an epithelium, realizing the stable terminal bifurcation ^27,52^. Luminal fluid is another potentially critical factor that can mechanically affect epithelial tissue deformation through shear stress and hydrostatic pressure ^53^. It has been previously reported that the manipulation of luminal pressure in murine developing lungs under ex vivo culture conditions altered branch formation ^36^. As oscillatory contractions of airway smooth muscles are likely to produce dynamic changes in the luminal pressure, it has been suggested that rhythmic luminal pressure would control the global synchronization of branching events in the murine lung ^37^, although it does not contribute to the tissue flattening during the terminal bifurcation.

The ERK-mediated curvature feedback mechanism proposed here advances the concept of mechanochemical regulation of the epithelial monolayer ^34,46,54,55^, in terms of taking the impact of tissue geometry into account. In various tissues, the epithelial cells collectively form their shape in the tissue scale and the constituent cells respond to the tissue curvature at different levels, including the transcription and cell activity levels ^56–59^. We believe that the proposed system works as a regulatory design principle underlying the mechanisms of self-organized tissue morphogenesis.

## Materials and Methods

### (I) Experiments

#### Animals

It has been previously reported that transgenic mice expressed hyBRET-ERK-NES, a FRET biosensor ^24–26^. The hyBRET-ERK-NES was comprised of cyan fluorescent protein (CFP) and yellow fluorescent protein (YFP), which acts as the donor and acceptor of FRET, respectively. Otherwise, we used ICR mice purchased from Japan SLC, Inc. We designated the midnight preceding the discovery of the vaginal plug as embryonic day 0.0 (E0.0), and all mice were sacrificed via cervical dislocation to minimize suffering. All animal experiments were approved by the local ethical committee for animal experimentation (MedKyo 18086, 19090, 20081, and 21043), and were performed in compliance with the guidelines for the care and use of laboratory animals at Kyoto University.

#### Antibodies, small molecules, and recombinant proteins

The followings were used: rat monoclonal anti-E-cadherin (Cell Signaling Technology, #3195, 1:100 dilution), rabbit polyclonal anti-phospho-myosin light chain (pMLC) (Abcam, #ab2480, 1:100 dilution), Alexa Fluor 546-conjugated goat anti-rat IgG (H+L) antibody (Thermo Fisher Scientific, #A11081, 1:1000), Alexa Fluor 647-conjugated goat anti-rabbit IgG (H+L) antibody (Abcam, #ab150079, or Thermo Fisher Scientific, #A21247, 1:1000), Blebbistatin (Merck Millipore, #203391), CK-666 (Sigma-Aldrich, #SML0006), Cytochalasin D (Merck Millipore, #250255), Jasplakinolide (Abcam, #ab141409), Latrunculin A (Cayman Chemical, #10010630), Nocodazole (Merck Millipore, #487928), PD0325901 (FUJIFILM Wako Pure Chemical Corporation, #162-25291), SMIFH2 (Sigma-Aldrich, #S4826), Y-27632 (Merck Millipore, #SCM075), FGF1 (R&D Systems, #232-FA), and FGF7 (R&D Systems, #251-KG).

#### Ex vivo culture

Dissected lung lobes were placed onto a hydrophilic polytetrafluoroethylene organ culture insert with a pore size of 0.4 µm (Merck Millipore, #PICM01250), which was preset in a 35 mm petri dish filled with 800 µL of culture medium. Samples were cultured in an air-liquid interface at 37°C under 5% CO_2_ conditions. To culture isolated epithelial tissues, dissected lobes were digested with 10 PU mL^-1^ of Dispase I (FUJIFILM Wako Pure Chemical Corporation, #386-02271) in phosphate buffered saline (PBS) for 15 min on ice, and immersed into fetal bovine serum (FBS) and kept on ice for 5 min, to stop the protease-mediated reaction. Then, samples were washed in PBS twice on ice, and the distal tips of epithelial tissues were manually excised using fine needles. Isolated epithelial tissues were cultured within growth factor-reduced Matrigel (Corning, #356231) and covered with 500 µL of culture medium added to each well of glass bottom dishes (Greiner, #627870) at 37°C under 5% CO_2_ conditions. We used FluoroBrite DMEM Media (Thermo Fischer Scientific, #A1896701) containing 1% GlutaMAX (Thermo Fischer Scientific, #35050061) and 0.5% FBS as the culture medium.

#### Whole-tissue immunofluorescence staining

Staining and optical clearing of dissected lung were performed according to previous studies ^60,61^. Briefly, samples were fixed with 4% paraformaldehyde (PFA) in PBS overnight at 4°C. For pMLC staining, samples were fixed with 2% trichloroacetic acid in PBS containing a 1% phosphatase inhibitor cocktail (1:100, Nacalai Tesque, #07575-51) for 15 min at 4°C. Then, samples were blocked by incubation in 10% normal goat serum (Abcam, #ab156046) diluted in 0.1% Triton X-100/PBS (PBT) for 3 h at 37°C. Samples were treated with primary antibodies overnight at 4°C, washed in 0.1% PBT, and subsequently treated with secondary antibodies conjugated to either Alexa Fluor 546 or Alexa Fluor 647 overnight at 4°C. For nuclear counterstaining, we used Hoechst33342 (5 µg ml^-1^, Dojindo Molecular Technologies, #H342) or DAPI (Dojindo Molecular Technologies, #D523-10, 1:200). The samples were mounted with 10 µL of 1% agarose gel onto a glass-based dish (Greiner Bio-One, #627871) for obtaining stable images. Then, samples were immersed with CUBIC-R+ (Tokyo Chemical Industry Co., # T3741) solution for 1 hour at 37°C or BABB solution (benzyl-alcohol and benzyl-benzoate, 1:2) for 30 min at 23°C.

#### F-actin and tubulin labeling

To visualize F-actin and microtubules, we used SiR-Actin and SiR-Tubulin, respectively ^62,63^, according to the manufacturer’s instructions (Cytoskeleton, Inc. #CY-SC001 and #CY-SC002). We added 1 µM of the far-red fluorogenic probes to the culture media, 1 hour before imaging. Although SiR-Actin was originally derived from Jasplakinolide, which activated ERK in isolated lung epithelial tissues in our experiments, ERK activation did not occur after treatment with SiR-Actin in isolated lung epithelial tissues.

#### Fluorescence labeling of FGF

To visualizes internalized FGFs, we used an amine-reactive pH-sensitive pHrodo iFL Red STP ester dye, according to the manufacturer’s instructions (Thermo Fischer Scientific, #P36014). For the imaging of FGF1, regardless of its localization inside or outside cells, we used the Alexa Fluor 488 Microscale Protein Labeling Kit, according to the manufacturer’s instructions (Thermo Fischer Scientific, #A30006).

#### Ethynyl deoxyuridine (EdU) assay

The lung epithelium was cultured within the Matrigel as described above for 3 hours with either 500 ng mL^-1^ FGF1 or 500 ng mL^-1^ FGF1 and 1 µM PD0325901 before the EdU incorporation to the lung epithelium. Next, 10 µM EdU dissolved into PBS was added to the culture media 20 min prior to the fixation with 4% PFA. Then, whole-tissue immunostaining of E-cadherin and nuclear counterstaining with DAPI were performed as described above. Then, EdU signal was detected using the Click-iT® EdU Imaging Kits (Thermo Fisher Scientific, #C10340) according to the manufacture’s instruction.

#### Microscopy

For live imaging, we used the incubator-integrated multiphoton fluorescence microscope system (LCV-MPE, Olympus) with a 25× water-immersion lens (NA=1.05, WD=2 mm, XLPLN25XWMP2, Olympus) or an inverted microscope (FV1200MPE-IX83, Olympus) with a 30× silicone-immersion lens (NA=1.05, WD=0.8 mm, UPLSAPO30XS, Olympus). The excitation wavelengths were set to 840 nm for the CFP of the ERK FRET biosensor, 930 nm for Alexa Fluor 488-labeled FGF1, and 1040 nm for the pHrodo iFL Red-labeled FGFs and the SiR probes (InSight DeepSee, Spectra-Physics). We used RDM690 IR cut filters (Olympus), and the DM505 and DM570 dichroic mirrors. We used BA460-500, BA495-540, BA520-560, BA575-630, and BA647/57 emission filters (Olympus) for CFP, Alexa Fluor 488, YFP or FRET, iFL Red probes, and SiR probes, respectively. For fixed samples, images were obtained using the Leica TCS SP8 confocal laser scanning platform equipped with the Leica HyD hybrid detector, using the 20× objective lens (NA=0.75, WD=680 μm, HC PL APO CS2, Leica) and 40× objective lens (NA = 1.3, WD = 240 μm, HC PL APO CS2, Leica).

#### FRET image analysis

The median filter of a 3 × 3 window was processed to remove shot noise, and the background signal was subtracted from each FRET and CFP channel. Then, the ratio of FRET intensity to CFP intensity was calculated using a custom-made MATLAB (MathWorks) script. In the scale bar, the color represents the FRET/CFP ratio and brightness represents the fluorescence intensity of the FRET channel.

#### Three-dimensional mapping

To determine the correlation between different quantities, such as ERK activity, F-actin signal, tissue curvature, and tissue thickness (Figure S1A), we obtained 3D stacked images of lung epithelial tissues with 1-µm-long z-intervals, and manually traced either the basal or apical side of the cells. Next, we generated triangle meshes on the traced surface using “Iso2Mesh,” a free mesh generation toolbox ^64^. Then, we calculated a discrete Gaussian curvature and a discrete curvature on each node of generated meshes using a custom-made code according to the definition proposed earlier ^65^. Finally, we obtained all values except those for the curvatures by averaging values around the nodes, and mapped them at each node. All processing was performed using MATLAB.

#### Curvature measurement in 2D

We used a spline curve-based method described elsewhere ^55^. Briefly, we manually traced the basal side of epithelial cells from images, and determined discrete sampling points (*x*_*i*_, *y*_*j*_) along traced curves at regular intervals. Upon fitting the 4 sampling points from *i*-1 to *i*+2 with a cubic spline function, the function *S*_*i*_ at an interval [*x*_*i*_, *x*_*i* + 1_] was denoted as *S*_*i*_ (*x*) = *a*_*i*_ (*x* − *x*_*i*_)^3^ + *b*_*i*_ (*x* − *x*_*i*_)^2^ + *c*_*i*_ (*x* − *x*_*i*_) + *d*_*i*_. Due to the definition of the curvature using the formula *κ*(*x*) = *S*″(1 + *S*′^2^) ^−3/2^, the curvature from the spline function was calculated as follows:

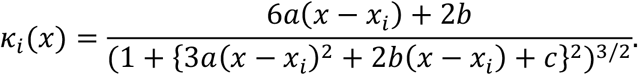

The convex/concave surface of the lumen was assigned a positive/negative value, namely *k*. We defined tissue thickness as the length from the apical to the basal edge, which is vertical to the traced curve at the sampling points.

#### Mechanical strain assay

We used a mechanical device (STREX, #STB-10) with a polydimethylsiloxane (PDMS) chamber (STREX, #STB-CH-04). To compress isolated epithelium, we placed isolated lung epithelial tissues in a 50% stretched state in the PDMS chamber, and added 1 mL of growth factor-reduced Matrigel (Corning, #356231), and performed gelation for 15 min at 37°C. Then, we added 2 mL of the culture medium supplemented with 500 ng mL^-1^ FGF1 into the chamber. We conducted experiments involving epithelial tissues under 2D culture conditions in the following manner. To obtain cell suspensions from primary tissues, lung tissues obtained via dissection at E12.5 were treated with 1 mg mL^-1^ Collagenase/Dispase (Sigma-Aldrich, #10269638001) in PBS for 50 min at 37°C while agitating the solution. Lung cells were then dispersed into the solution by gentle pipetting, and the suspended solution was filtered using a 40 µm pore size mesh (Corning, #352340). After centrifugation at 200G for 5 min at 25°C, the cell pellet was resuspended in a culture medium containing 10 µM Y-27632 and 10% KnockOut Serum Replacement SR (Thermo Fischer Scientific, # 10828010) and the solution was placed in a PDMS chamber pretreated with a collagen coating (Cellmatrix Type I-C, Nitta Gelatin, #631-00771). After 2 days, the culture medium was exchanged with a medium not containing Y-27632 and KSR, 1 hour before imaging. Notably, the PDMS chamber gradually droops by becoming attached to the silicone oil used for objective lenses, because of which the focus has to be adjusted manually every time an image needs to be obtained.

#### Osmolyte-based compression assay

The osmolyte-based tissue compression was performed according to the previous studies ^30,66^. The lung epithelium was cultured within the Matrigel for 3 hours with 500 ng mL^-1^ FGF1 before the dextran administration. We used the dextran of 1500-2800 kDa in average molecular weight (Merck, #D5376) to prevent penetrating into the Matrigel. The culture medium was partially replaced into the dextran solution dissolved in the medium to reach the final concentrations for the compression. We converted the dextran concentration to the pressure based on the function of pressure versus dextran concentration ^66^: 30 mg mL^-1^ for 2 kPa, and 55 mg mL^-1^ for 5 kPa.

#### Statistical hypothesis testing

The number of cells or regions of interest analyzed (n) and the number of biological replicates (N) are indicated in the figure legends. No particular statistical method was used to predetermine the sample size. A minimum of N=3 independent experiments were performed, based on previous studies in the field. No inclusion/exclusion criteria were used and all analyzed samples were included in the analysis. No randomization was performed. Statistical tests, sample sizes, test statistics, and *P*-values have been described in the main text. We considered P-values < 0.05 to be statistically significant in two-tailed tests, and classified them into the following 4 categories: * (p<0.05), ** (p<0.01), *** (p<0.001), and n.s. (not significant, i.e., p ≥ 0.05).

#### Software

We used MATLAB (MathWorks) and Image J (National Institute of Health) software for digital image processing. For graphics, MATLAB (MathWorks), Imaris (Bitplane), and Image J (National Institute of Health) were used. MATLAB (MathWorks) was used for statistical analysis.

#### Graph

In the boxplot, the central mark indicates the median, and the bottom and top edges of the box indicate the 25^th^ and 75^th^ percentiles, respectively. The whiskers extend to the most extreme data points not considered to be outliers, and the outliers are plotted individually using the ‘+’ symbol. All the graphs were drawn using MATLAB.

### (II) Model analysis

#### (IIa) Apical length vs cell height at equilibrium

Here, we consider cells in a flat epithelial sheet to be identical, and assume that the basal edge length is almost the same as the apical edge length for the simplicity of analysis. Then, potential energy can be defined as follows:

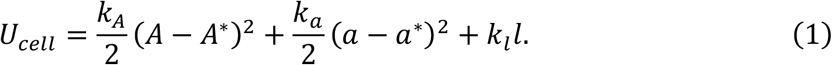

The first term represents the cell size constraint and its coefficient *k*_*A*_, current cell area *A*, and the target cell area *A*^∗^. The second and third terms each represent the regulation of cell edge length at the apical and lateral side of cells *a* and *l* with controlling parameters *k*_*a*_ and *k*_*l*_. We set the second term as a function that converges to the variable target apical length *a*^∗^, as explained later. Since the cell size was maintained even after a change in cell shape, caused by the inhibitor assay, we set *A* = *A*^∗^. Then, upon considering that *A* = *rh*, we can redefine equation 1 as follows:

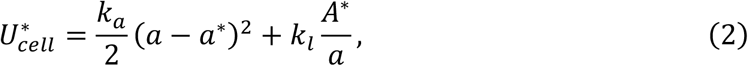

Now, we have two variables, *a* and *a*^∗^, and obtained equilibrium values at the minimum potential energy value, i.e., 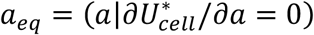 and 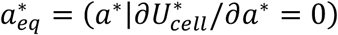 each, as 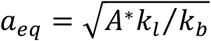 and 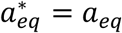. From these equations, it was noted that 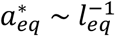, which indicates the existence of an inverse relationship between the apical length and cell height at equilibrium. The analysis in this section is based on that of a previous study ^67^.

#### (IIb) Apical length vs epithelial sheet curvature

The analysis in this section is based on that of a previous study ^68^. Here, we considered the geometrical aspects of an epithelial monolayer, composed of smoothly-connected identical tetragons along a line. The curvature *κ* at the midplane of the epithelial sheet is represented as

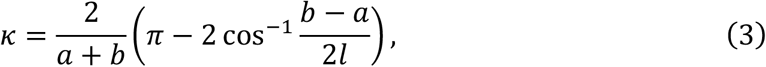

where *a, b*, and *l* denote the apical, basal, and lateral edge lengths, respectively. Then, the curvature with respect to the apical edge length is determined as follows:

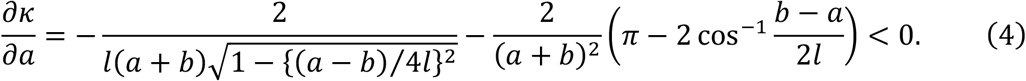

This indicates that the sheet curvature becomes small with an increase in the apical edge length.

#### (IIc) Mathematical modeling for multicellular dynamics

We modeled an epithelial monolayer using the vertex model framework ^32–34^. In our model, a single cell is represented as a polygon with four vertices and edges, which are shared by neighboring cells. The dynamics of vertex position **r**_*i*_, where *i* is an index of the vertex, obeys the equation of motion, based on the principle of least potential energy *U*, with a drag coefficient *η* and luminal pressure *p* as follows:

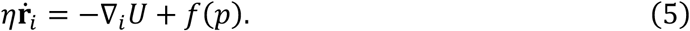

To determine the potential energy in the first term, we defined a minimal expression:

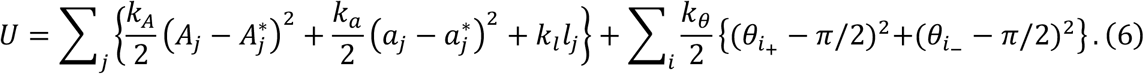

The first term represents the cell size constraint with its coefficient *k*_*A*_, current cell area *A*_*j*_, and the target cell area 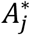. The second and third terms each represent the regulation of the cell edge length at the apical and the lateral sides of cells *a*_*j*_ and *l*_*j*_, with controlling parameters *k*_*a*_ and *k*_*l*_. As the apical edge is controlled by the ERK activation level, we set the second term as a function that converges to the variable target apical length 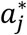; this has been explained later. The fourth and fifth terms represent the bending energy of cells at the apical and the basal sides, attributed to each vertex *i*; *k*_*θ*_ denotes the bending rigidity, and 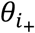 and 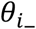 are angles at the vertex *i* across the cell edge.

The second term of Equation 5 represents the mechanical effect of luminal pressure ^36,37,53^ on the apical edge of epithelial cells. The function *f* was defined as

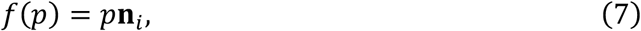

where *p* is strength of pressure, and **n**_*i*_ is the normal vector at the apical vertex *i*. Note that this function is effective only at vertices in the apical sides of cells.

To implement cell proliferation, we ensured that each cell was set with the cell timer *τ*_*j*_, and all cells had the same cell cycle length *τ*_*div*_. We assumed that cell cycle progression depended on the ERK activity, namely the curvature in the model; i.e., the cell cycle progresses when the cell curvature is not less than zero (*k*_*j*_ ≥ 0), and the cell cycle pauses when the cell curvature is less than zero (*k*_*j*_ < 0), according to the previous report ^38^ and cell proliferation assay (Figure S6). This is also attributable to the fact that cell proliferation in the developing murine lung requires ERK activation, which can be observed in the distal tip ^23,69^. Cell division occurs and cells with almost equal sizes are produced when the cell timer reaches the cell cycle length, *τ*_*j*_ = *τ*_*div*_. Once cell division occurs, the *τ*_*j*_ value in one of the daughter cells resets to zero and that in another is set at a value chosen stochastically from 0 to 0.05*τ*_*div*_ with a uniform distribution, to avoid perfect synchronization between neighboring cells.

In our model, the target apical edge length depends on the cytoskeletal level *c*, observed with a level of F-actin accumulation, as follows:

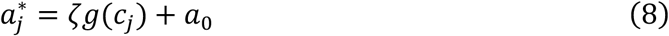

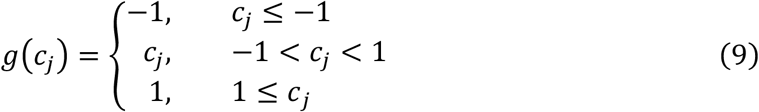

Here, *ζ* denotes the maximum length after edge extension, and *a*_0_ is the baseline for the apical edge length. The saturation function *g* represents the physical limit for the change in edge length. We modeled the cytoskeletal level *c* as follows:

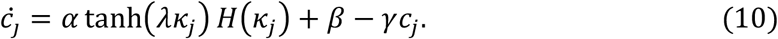

The first term *ċ*_*j*_ represents the curvature-dependent contribution of FGF-ERK, along with the magnitude parameter *α* and the sensitivity parameter *λ*, to the curvature *κ. H* is the step function of the curvature, i.e., *H*=1 for *κ* ≥ 0 and *H*=0 for *κ* < 0. The curvature *k*_*j*_ at cell *j* is calculated using the spline function-based method described above. As the cytoskeleton level controls the apical target length observed with an increase in the monotonic function, as shown in Equation 9, curvature-dependent apical extension/shrinkage is represented as a positive/negative *α* value. The second term represents the curvature-independent contribution of FGF-ERK, along with the parameter β. The third term represents the decay of cytoskeletons, including F-actin depolymerization and severing. The rate *γ* controls the characteristic decay time; the inverse value of *γ* denotes the retention time for cell curvature memory. The level of curvature-dependent FGF-ERK activation was determined in Equation 10 because of its rapid response.

#### (IId) Numerical simulations

We considered a manually traced line along the basal sides of cells in the image shown in Figure 1 as the initial shape for the simulation, when the diameter of the distal tip *D* was in the measured range. Upon an analysis of the tissue morphology, the number of cells was determined according to the measured cell size, i.e., the average cell width and cell height, which were 5 µm and 20 µm, respectively. The boundary in the proximal side was fixed in the simulations shown in Figures 6B and 6C. For the simulations in Figure 6D–I, we used the symmetrical image created based on the Figure 6A. Ordinary differential equations were numerically solved using the forward Euler method, with a time step of 0.01, using MATLAB (MathWorks). The simulations were terminated when the number of cells was more than 300.

Standard parameters were set as follows: *η* = 1, *k*_*A*_ = 0.01, *k*_*a*_ = 1, *k*_*l*_ = 0.2, *k*_*θ*_ = 20, *τ*_*div*_ = 300, *α* = 1, *β* = 0, *γ* = 0.001, *ζ* = 3 μ*m, λ* = 100, *p* = 0.05, and *D* = 70 μ*m*, unless otherwise noted. The *k*_*A*_, *k*_*a*_, *k*_*l*_, *k*_*θ*_, *λ*, and *p* values were determined by numerical analyses using several criteria, under which the simulated curvature imitates the experimental observations and became insensitive to the change in the values, as shown in Figure S5. The maximum length of edge extension, i.e., the *ζ* value, was determined from our measurements. 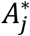 and *a*_0_ values were set according to the initial conditions, e.g., 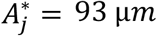 and *a*_0_ = 4 μ*m* when *D* = 70 μ*m*.

## Acknowledgements

This work was supported by the JST PRESTO grant JPMJPR1949, by the JSPS KAKENHI 19H00993 and 21H05290, and by the Medical Research Support Center of Kyoto University. We would like to thank Yu Kurata and Akane Kusumi for technical assistance, Edouard Hannezo, Naoya Hino, Mamoru Ishii, and Takuya Yoshida for fruitful discussion.

## Data Availability

Source data are provided with this paper.

## Code Availability

Relevant codes for this research work are stored in GitHub: https://github.com/tsuyoshihirashima/vertex-lung

## Author Contributions

Conceptualization, T.H.; Methodology, T.H., M.M.; Software, T.H.; Validation, T.H.; Formal analysis, T.H.; Investigation, T.H.; Resources, T.H., M.M.; Data curation, T.H.; Writing - original draft, T.H.; Writing - review & editing, T.H., M.M.; Visualization, T.H.; Supervision, T.H.; Project administration, T.H., M.M.; Funding acquisition, T.H., M.M.

## Supplementary figure legends

**Figure S1** 3D mapping of ERK activity and tissue geometry

(A) Mapping procedure for linking different quantities, including the ERK activity, epithelial sheet curvature, and sheet thickness. Details of the procedure are described in the Materials and Methods section.

(B) The relationship between the ERK activity and discrete mean curvature. ρ=0.61±0.02, where ρ is Spearman’s rank correlation coefficient. Data represent mean and SD values. n=1495, N=3.

(C) Time-lapse images of ERK activity (upper) and bright field (lower) during terminal bifurcation of an epithelium in the intact lung lobe under the explant culture condition. Yellow dotted line represents the basal side of the epithelial tissue. Green arrows represent the ERK activation. Scale bars, 50 µm.

**Figure S2** FGF1 endocytosis from the basal side of an isolated lung epithelium

(A) Time-lapse images of the bright field (upper), FGF1-pHrodo (middle), and FGF1-Alexa488 (lower). Red and blue circles in the bright field images indicate the tip apex and the cleft bottom, respectively. The time origin indicates the time of administration of labeled FGF1. FGF1 is distributed only in the Matrigel, not in the lumen, indicating that FGF1 internalization-based endocytosis occurs from the basal side of the epithelium. Notably, FGF1 distribution is almost homogeneous in the Matrigel (lower), indicating that no clear gradient was formed near the epithelium, even after ligand internalization. Scale bar, 50 µm.

(B) FGF1-pHrodo intensity at the tip apex (red) and the cleft bottom (blue) over time. The response time of FGF1-pHrodo at the tip apex is 86 min. N=3.

(C) FGF1-Alexa488 intensity in the Matrigel over time. The response time of FGF1-Alexa488 in the Matrigel is 26 min. N=3.

**Figure S3** Dynamics of ERK activity in the growing lung epithelium with FGF7

Time-lapse images of ERK activity (upper) and bright field (lower) in the case of 100 ng mL^- 1^ FGF7 treatment to the isolated lung epithelial tissue. Time origin is at 4 hours after the FGF1 treatment. Scale bar, 30 µm.

**Figure S4** Morphological changes owing to chemical perturbations and cytoskeletal architecture

(A) Morphological change in the isolated epithelium after treatment with 1 µM PD0325901. Colors indicate the ERK activity level. Scale bar, 30 µm.

(B) Quantification of cell morphological parameters: cell height (left), width (middle), and size (right). n=40, N=4. Welch’s t test: p<0.001, p<0.001, p<0.001 (cell height: Control-Jasp, Control-LatA10min, Control-LatA20min), p<0.001, p=0.019, p=0.46 (cell width: Control-Jasp, Control-LatA10min, Control-LatA20min), and p=0.663, p=0.675, p<0.001 (cell size: Control-Jasp, Control-LatA10min, Control-LatA20min)

(C) Isolated lung epithelium stained with SiR-Actin. Scale bar, 30 µm.

(D, E) Morphological change of epithelium (D) and measured relative change in cell height (E) after treatment with the mixture of 1 µM Latrunculin A and 1 µM Nocodazole. N=6. Scale bar, 30 µm.

(F) Isolated lung epithelium stained with SiR-Tubulin. Scale bar, 30 µm.

(G–G”) Immunofluorescence images of cells interacting with anti-E-cadherin (blue), anti-phospho myosin light chain (pMLC, magenta) and the Hoechst (white) nuclear counterstain. Notably, the pMLC signal was significantly weaker at the tip of the epithelium than in smooth muscle cells around the stalk. In the magnified view (G”), the pMLC signal was clear at the microtubule organizing centers (arrowhead), and was localized at the apical side of cells despite its weak intensity. Scale bars, 100 µm (G, G’) and 20 µm (G”).

**Figure S5** Simulation of the mathematical model

(A) Parameter dependency of the monolayer curvature in the simulation.

(B) Relative cell height, normalized by one at *α*=0, as a function of *α*. Representative morphologies of the virtual monolayer sheet are shown in the upper image.

(C) Tissue curvature as a function of luminal pressure *p*. Representative morphologies of virtual monolayer tissues are shown in the upper image. N=5.

**Figure S6** EdU-based proliferation assay

Fluorescence labeling images describing EdU (magenta), nuclear DAPI counterstaining (blue), and anti-E-cadherin (green) in the isolated epithelium, treated with 500 ng mL^-1^ FGF1 (upper) and the mixture of 500 ng mL^-1^ FGF1 and 1 µM PD0325901 (MEK inhibitor) (lower), respectively. EdU signal was detected mostly in the distal tips in the FGF1 treatment, while it was undetected in the inhibition of ERK activity. Scale bars, 20 µm.

## Movie legends

**Movie 1 Volumetric imaging of ERK activity in the murine lung**

The 3D ERK activity map of a murine lung dissected at E12.5. Colors represent the ERK activity and the color scale corresponds to that in Figure 1B. Scale bar, 50 µm.

**Movie 2 Dynamics of ERK activity in the developing lung under the explant culture condition**

Time-lapse video of bright field (left) and ERK activity (right) during the developing lung. The color scale corresponds to that in Figure S1C. Scale bar, 50 µm.

**Movie 3 Endocytosis of FGF1 in the isolated lung epithelium**

Time-lapse video of bright field (left), FGF1-Alexa488 (middle), and FGF1-pHrodo (right) in the isolated lung epithelium embedded within the Matrigel. The color scales correspond to those in Figure S2A. The time origin indicates the time of administration. Scale bars, 50 µm.

**Movie 4 Dynamics of ERK activity in the growing lung epithelium with FGF7**

Time-lapse video of bright field (left) and ERK activity (right) during the morphological change of an isolated lung epithelium treated with 100 ng mL^-1^ FGF7. The color scale corresponds to that in Figure S3. Scale bars, 30 µm.

**Movie 5 Dynamics of ERK activity in the growing lung epithelium with FGF1**

Time-lapse video of bright field (left) and ERK activity (right) during the terminal bifurcation of an isolated lung epithelium treated with 500 ng mL^-1^ FGF1. The color scale corresponds to that in Figure 2D. Scale bars, 30 µm.

**Movie 6 ERK activity response in the isolated lung epithelium to the uniaxial compression parallel to the distal-proximal axis**

Time-lapse video of ERK activity at the tip of the isolated lung epithelium against compression parallel to the distal-proximal axis. The color scale corresponds to that in Figure 3B. The time origin indicates the timing of compression. Scale bar, 30 µm.

**Movie 7 ERK activity response of the isolated lung epithelium to the uniaxial compression vertical to the distal-proximal axis**

Time-lapse video of ERK activity at the tip of an isolated lung epithelium against the compression vertical to the distal-proximal axis. The color scale corresponds to that in Figure 3C. The time origin indicates the timing of compression. Scale bar, 30 µm.

**Movie 8 Morphological response of the isolate epithelium to the osmolyte-based global compression**

Time-lapse video of bright field for the isolated lung epithelium in the compression assay of 5 kPa. Time origin indicates the timing of dextran administration. Scale bar, 50 µm.

**Movie 9 Morphological change in the isolated lung epithelium by ERK inactivation**

Time-lapse video of ERK activity at the tip of the isolated lung epithelium after treatment with 1 µM PD0325901. The color scale corresponds to that in Figure S4A. Time origin indicates the timing of inhibitor administration. Scale bar, 30 µm.

**Movie 10 Response of ERK activity and F-actin accumulation to acute simulation with FGF1**

Time-lapse video of ERK activity (left) and SiR-Actin signal (right) at the tip of the isolated lung epithelium after treatment with 500 ng mL^-1^ FGF1. The color scales correspond to those in Figure 5A. The time origin indicates the timing of FGF1 administration. Scale bars, 30 µm.

**Movie 11 Simulation for the morphogenesis of the proliferating epithelial monolayer in the regime of curvature-dependent apical extension**

Colors represent the sheet curvature. The color scale corresponds to that in Figure 6E. Scale interval of the graph is 20 µm.

**Movie 12 Simulations of virtual epithelial monolayer with the short (γ=1), medium (γ=10**^**-3**^**), and long (γ=10**^**-3**^**) retention time for curvature memory**

Colors represent the cellular F-actin levels. The color scale corresponds to that in Figure 6H. Scale interval of the graph is 20 µm.

