## Supplementary figures and images for "ERK-mediated Curvature Feedback Regulates Branching Morphogenesis in Lung Epithelial Tissue"

### Supplemental Figures

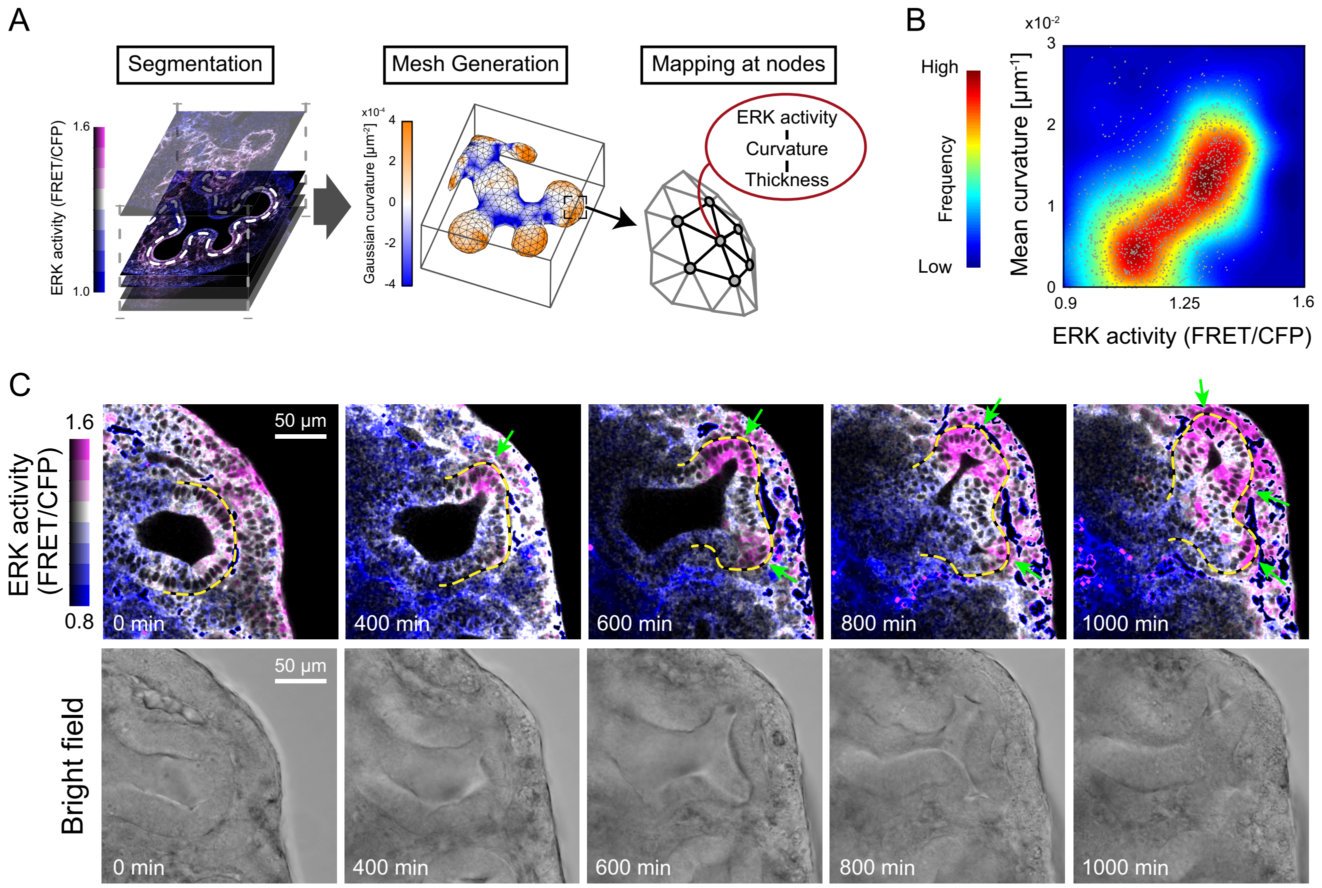

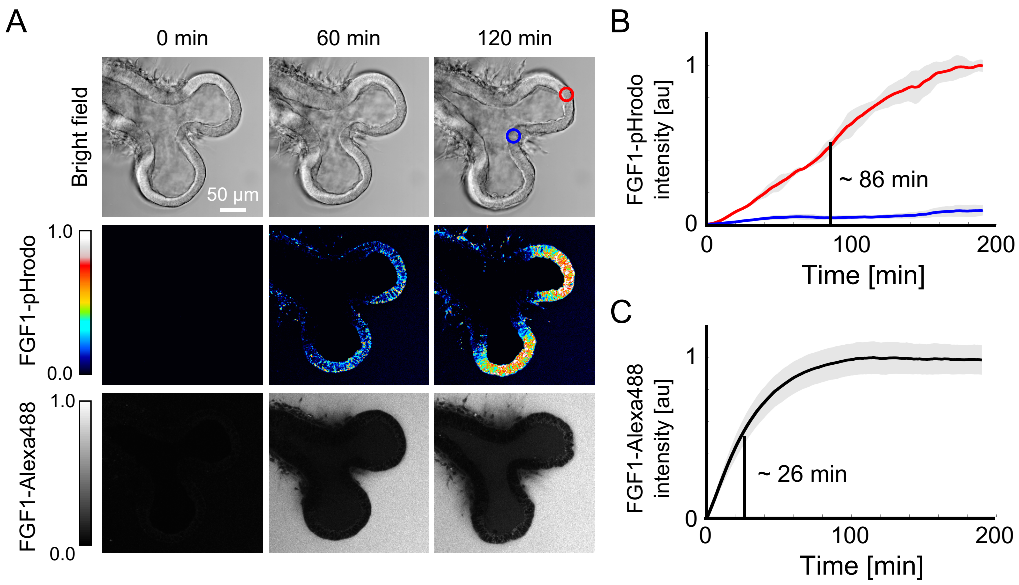

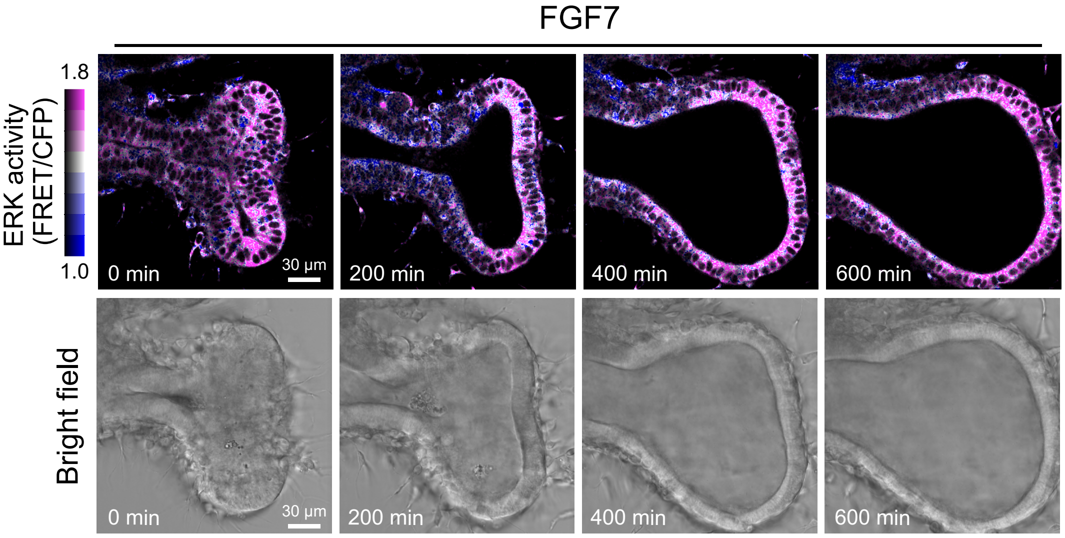

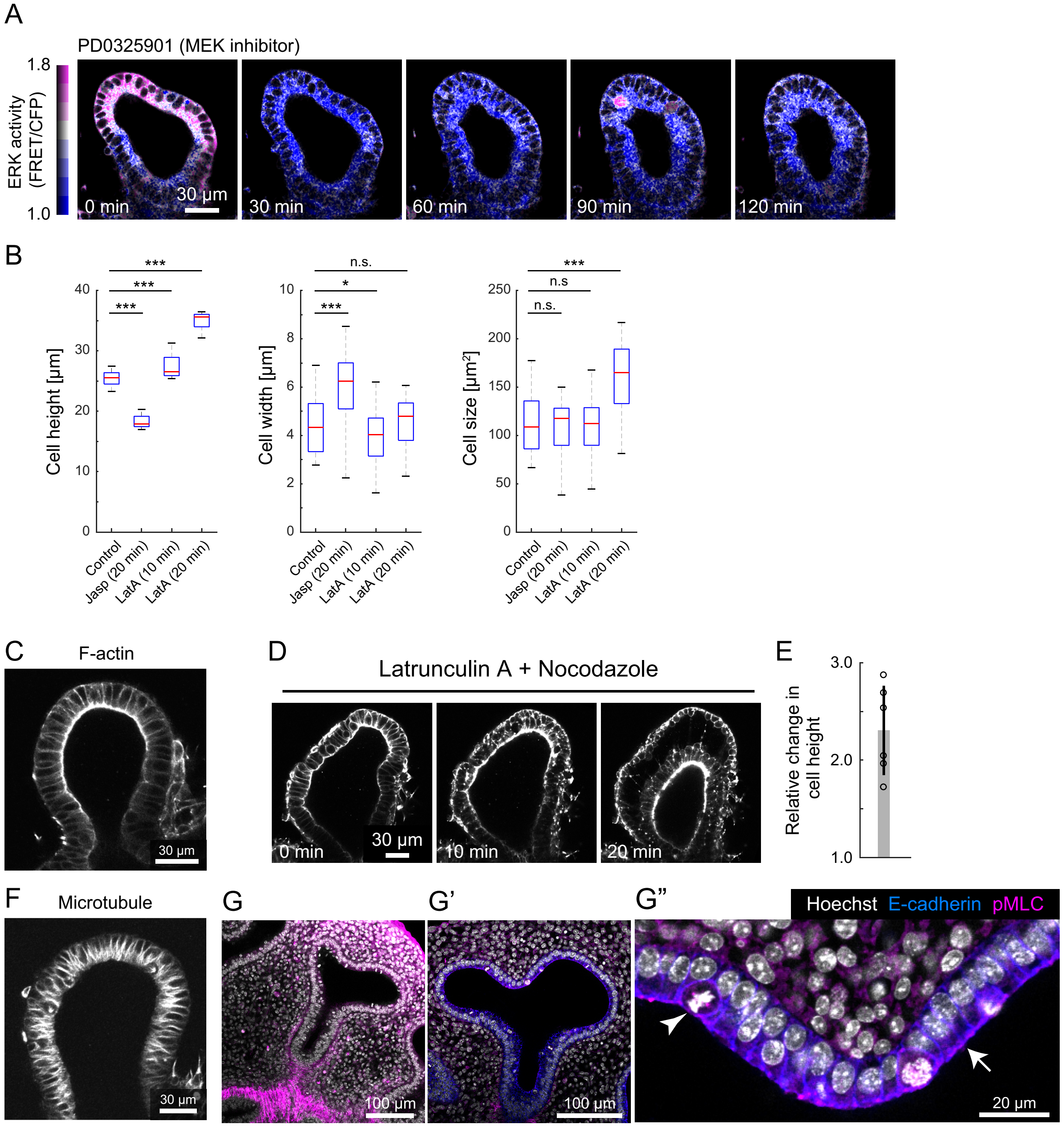

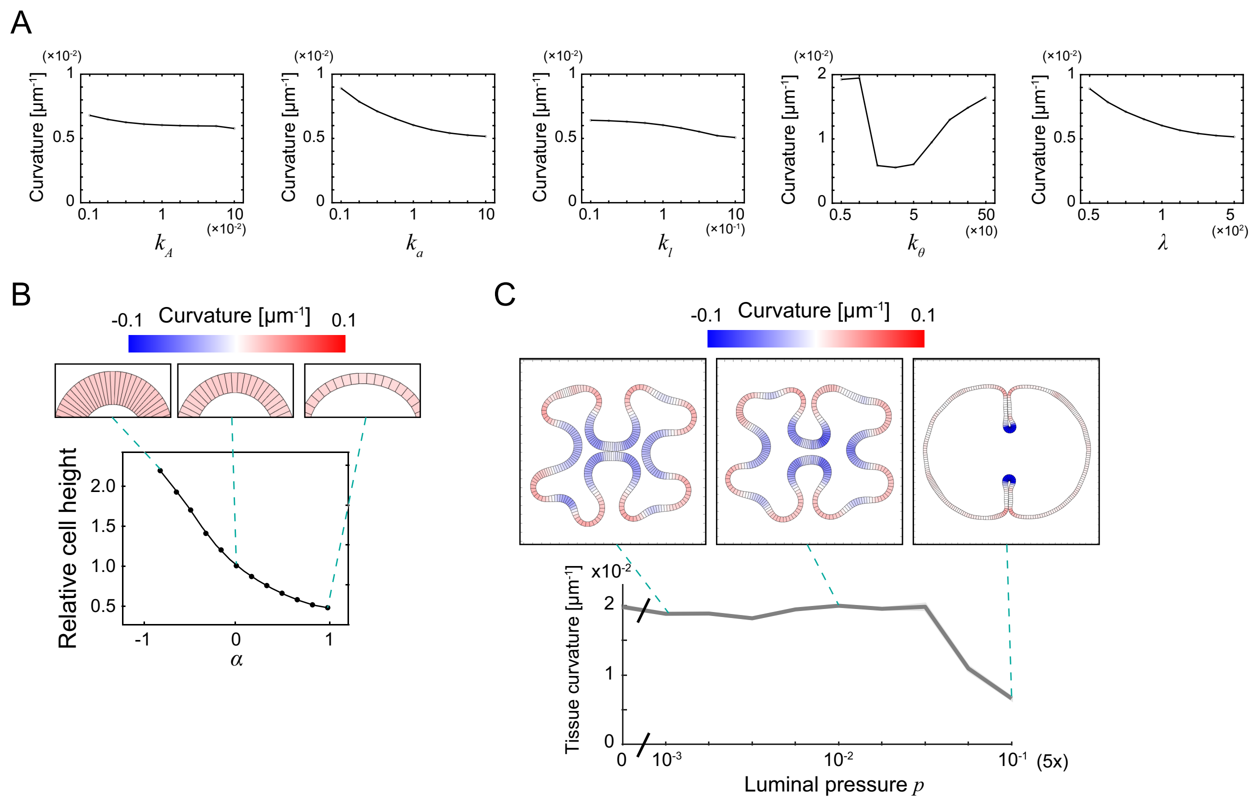

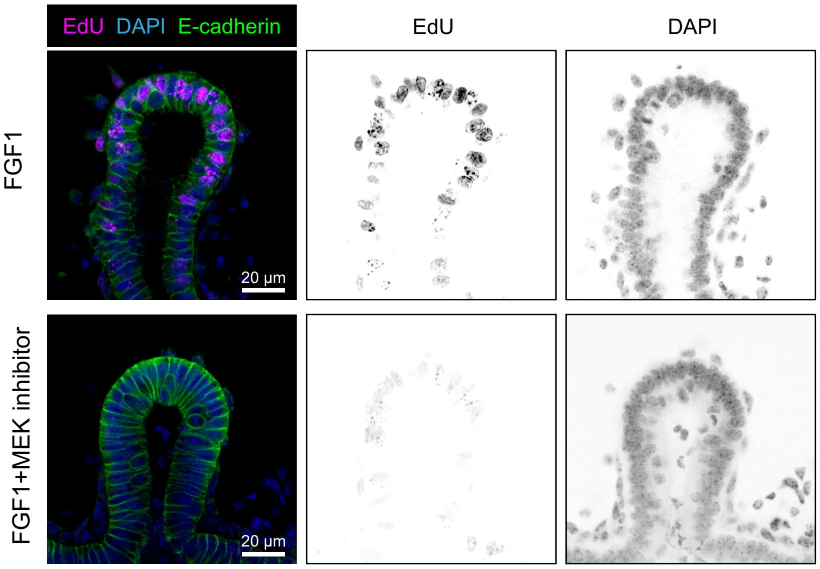
